# Remyelination alters the pattern of myelin in the cerebral cortex

**DOI:** 10.1101/2020.03.09.983569

**Authors:** Jennifer Orthmann-Murphy, Cody L. Call, Gian Carlo Molina-Castro, Yu Chen Hsieh, Matthew N. Rasband, Peter A. Calabresi, Dwight E. Bergles

## Abstract

Destruction of oligodendrocytes and myelin sheaths in cortical gray matter profoundly alters neural activity and is associated with cognitive disability in multiple sclerosis (MS). Myelin can be restored by regenerating oligodendrocytes from resident progenitors; however, it is not known whether regeneration restores the complex myelination patterns in cortical circuits. Here we performed time lapse *in vivo* two photon imaging in somatosensory cortex of adult mice to define the kinetics and specificity of myelin regeneration after acute oligodendrocyte ablation. These longitudinal studies revealed that the pattern of myelination in cortex changed dramatically after regeneration, as new oligodendrocytes were formed in different locations and new sheaths were often established along axon segments previously lacking myelin. Despite the dramatic increase in axonal territory available, oligodendrogenesis was persistently impaired in deeper cortical layers that experienced higher gliosis. The repeated reorganization of myelin patterns in MS may alter circuit function and contribute to cognitive decline.

## INTRODUCTION

Oligodendrocytes form concentric sheets of membrane around axons that enhance the speed of action potential propagation, provide metabolic support and control neuronal excitability through ion homeostasis. Consequently, loss of oligodendrocytes and myelin can alter the firing behavior of neurons and impair their survival, leading to profound disability in diseases such as multiple sclerosis (MS), in which the immune system inappropriately targets myelin for destruction. In both relapsing-remitting forms of MS and the cuprizone model of demyelination in mouse, the CNS is capable of spontaneous remyelination through mobilization of oligodendrocyte precursor cells (OPCs) (Baxi et al., 2017; Chang, Nishiyama, Peterson, Prineas, & Trapp, 2000; Chang et al., 2012), which remain abundant in both gray and white matter throughout adulthood (Dimou, Simon, Kirchhoff, Takebayashi, & Götz, 2008; Hughes, Kang, Fukaya, & Bergles, 2013; Young et al., 2013). The highly ordered structure of myelin in white matter tracts has enabled *in vivo* longitudinal tracking of inflammatory demyelinating lesions using magnetic resonance imaging (MRI); however, due to the low spatial resolution of standard MRI sequences (Oh et al., 2019) and the indirect nature of MR methods used to assess the integrity of myelin (Walhovd, Johansen-Berg, & Káradóttir, 2014), our knowledge about the dynamics of OPC recruitment, oligodendrogenesis and remyelination within specific neural circuits remains limited.

*In vivo* studies of remyelination have focused primarily on white matter, where assessments of myelin are aided by the high density and symmetrical alignment of myelin sheaths; however, postmortem histological analysis (Kidd et al., 1999; Kutzelnigg, 2005; Lucchinetti et al., 2011; Peterson, Bö, Mörk, Chang, & Trapp, 2001) and new *in vivo* MRI and PET imaging methods (Beck et al., 2018; Filippi et al., 2014; Herranz et al., 2019; Kilsdonk et al., 2013; R. Magliozzi, Reynolds, & Calabrese, 2018) indicate that demyelination is also prevalent in cortical gray matter of MS patients. Cortical lesion load correlates with signs of physical and cognitive disability, including cognitive impairment, fatigue and memory loss (Calabrese et al., 2012; Nielsen et al., 2013). Nevertheless, much less is known about the capacity for repair of myelin in cortical circuits. Defining how gray matter lesions are resolved *in vivo* is critical for both MS prognosis and the development of new therapies to promote remyelination.

Unlike white matter, myelination patterns in the cerebral cortex are highly variable, with sheath content varying between cortical regions, among different types of neurons and even along the length of individual axons (Micheva et al., 2016; Tomassy et al., 2014). Despite this evidence of discontinuous myelination, recent *in vivo* imaging studies indicate that both oligodendrocytes and individual myelin sheaths are remarkably stable in the adult brain (Hill, Li, & Grutzendler, 2018; Hughes, Orthmann-Murphy, Langseth, & Bergles, 2018), suggesting that maintaining precise sheath placement is important for cortical function. However, the complex arrangement of cortical myelin presents significant challenges for repair and it is unknown whether precise myelination patterns in the cortex are restored following a demyelinating event.

*In vivo* two photon fluorescence microscopy allows visualization of oligodendrogenesis and myelin sheath formation in mammalian circuits at high resolution, providing the means to define both the dynamics and specificity of regeneration (Hughes et al., 2018). Here, we used this high-resolution imaging method to define the extent of oligodendrocyte regeneration and the specificity of myelin replacement within the adult somatosensory cortex after demyelination. Unexpectedly, our studies indicate that oligodendrocytes are regenerated in locations distinct from those occupied before injury. Despite the additional available axonal territory for myelination, regenerated oligodendrocytes had a similar size and structure; as a result, only a fraction of prior sheaths were replaced and many new sheaths were formed on previously unmyelinated regions of axons. Conversely, in regions of high territory overlap, new oligodendrocytes often formed sheaths along the same segment of specific axons, demonstrating that precise repair is possible. Together, these *in vivo* findings indicate that regeneration of oligodendrocytes in the cortex creates a new pattern of myelination, with important implications for the restoration of sensory processing and cognition.

## RESULTS

### Inefficient regeneration of oligodendrocytes in cortical gray matter

To define the dynamics of oligodendrocyte regeneration and axonal remyelination in the cerebral cortex, we performed longitudinal two photon imaging through a cranial window placed over the barrel field of the somatosensory cortex in transgenic mice that express EGFP under control of the promoter/enhancer for myelin-associated oligodendrocyte basic protein (*Mobp-EGFP*) mice (Hughes et al., 2018) (Figure 1A). In these mice, complete oligodendrocyte morphologies could be resolved *in vivo*, including cytoplasmic processes and individual myelin internodes within the upper layers of the cortex. In these regions, oligodendrocytes ensheath a select group of axons, including long-range axonal projections oriented horizontally to the pia (Bock et al., 2011) (Figure 1B), and in deeper layers, vertically-oriented axons belonging to both local cortical neurons and long-range projections (Figure 1A,C). To induce demyelination, young adult *Mobp-EGFP* mice (age 8-12 weeks) were fed chow mixed with 0.2% cuprizone, a copper chelator that induces robust fragmentation and apoptosis of oligodendrocytes (Vega-Riquer, Mendez-Victoriano, Morales-Luckie, & Gonzalez-Perez, 2019) (Supplementary Figure 1), and multiple volumes (425 μm × 425 μm × 550 μm) corresponding to layers I–IV were imaged repeatedly prior to injury, during demyelination and through recovery for up to 12 weeks (Figure 1D, Supplementary Video 1).

**Figure 1:**
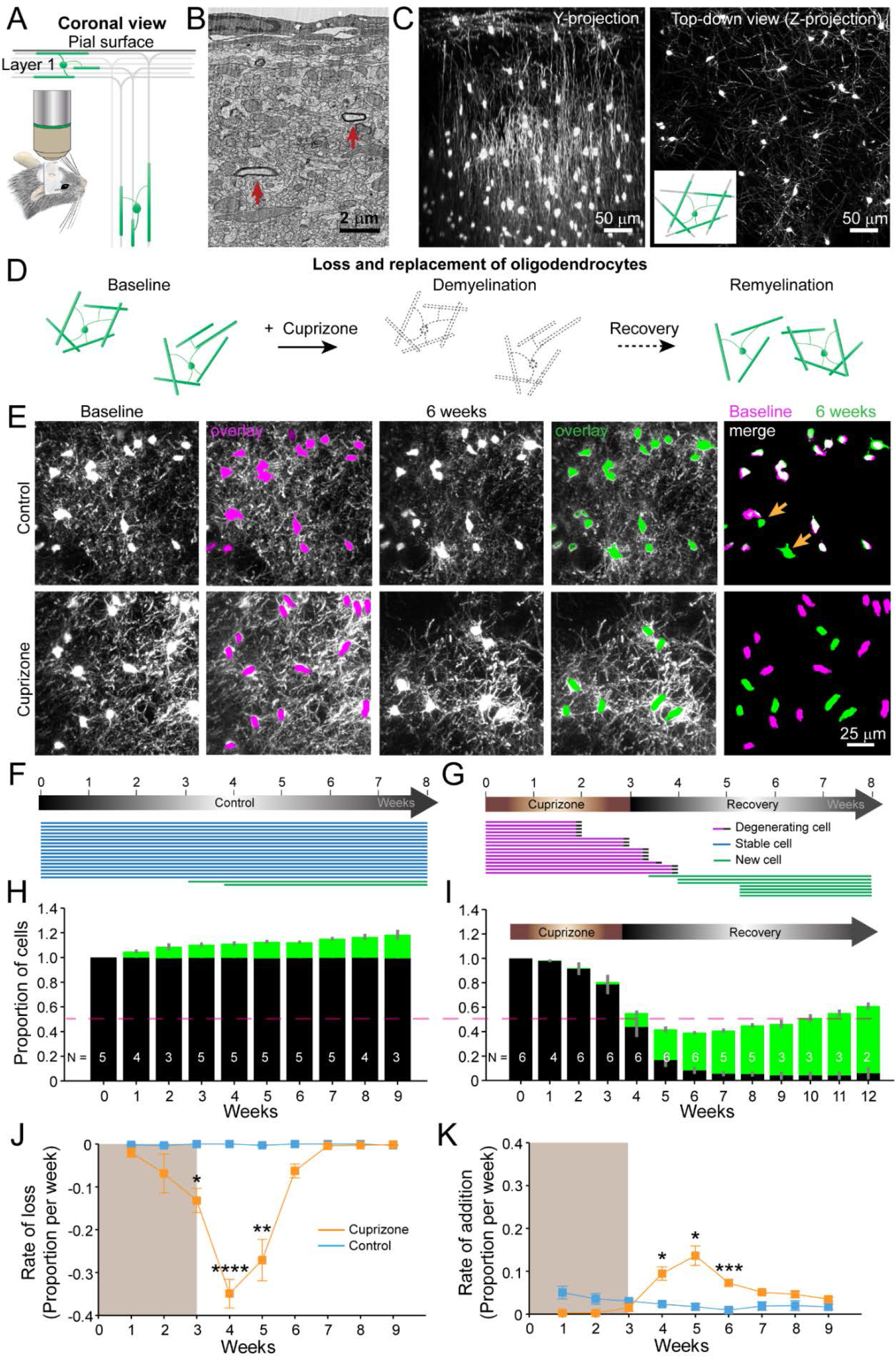
An *in vivo* platform to monitor loss and replacement of oligodendrocytes in the upper cortical layers. A: *In vivo* two photon microscopy through chronic cranial windows over the somatosensory cortex of *Mobp-EGFP* mice (coronal view), showing myelinated fibers in cortical layer I parallel to pial surface and in deeper layers oriented perpendicularly. B: Electron micrograph reconstruction of adult mouse visual cortex (from (Bock et al., 2011) illustrating low density of myelinated fibers (arrows) in the upper layers of cortex. C: Maximum intensity *y*-projection (coronal view, 425 μm × 150 μm × 550 μm) and *z*-projection (top-down view, 425 μm × 425 μm × 100 μm) example regions from *Mobp-EGFP* mice with chronic cranial windows. D: Schematic illustrating longitudinal course of loss (demyelination) and replacement (remyelination) of cortical oligodendrocytes. E: Examples of maximum intensity projection images of the same region (156 μm × 156 μm × 84 μm) imaged repeatedly from an adult sham- (control, top row) or a cuprizone-treated (bottom row) mouse are shown with overlay of cell bodies from baseline (magenta) and after 6 weeks (green). Merge of baseline and 6 week overlays show where new cells are added to the region (arrows). F-G: Individual cells (represented by magenta, blue or green lines) were tracked longitudinally in somatosensory cortex from mice fed control (F; from region in top row of E) or cuprizone diet (G; from region in bottom row of E). H-K: The same cortical volume (425 μm × 425 μm × 300 μm) was imaged repeatedly in mice given either control or cuprizone diet, and individual cells present at baseline (black) or formed at later time points (green) were tracked over time. Shown are the average cell counts depicted as a proportion of baseline number of cells, (H, N = 5 control mice; I, N = 6 cuprizone mice, I; number of mice imaged at each time point indicated). J-K: The average rate of loss (J) or addition (K) of oligodendrocytes per week in control-treated (blue) v. cuprizone-treated mice (orange) relative to the baseline population of oligodendrocytes. Treatment with sham or cuprizone-supplemented chow denoted by shaded background. In cuprizone-treated mice, there was a higher rate of oligodendrocyte loss over weeks 3-5 (@ 3 weeks, *p* = 0.02014; @ 4 weeks, p=0.0000488; @ 5 weeks, *p* = 0.0066) and addition of new cells between 4-6 weeks (@ 4 weeks, *p* = 0.0280; @ 5 weeks, *p* = 0.0121; @ 6 weeks, *p* = 0.000530) compared to control. Data is presented as means with standard error of the mean bars; data are compared with N-way ANOVA test with Bonferroni correction for multiple comparisons.

Oligodendrocytes and individual myelin sheaths are extraordinarily stable in the adult brain (Hill et al., 2018; Hughes et al., 2018; Yeung et al., 2014); however, the amount of myelin within these circuits is not static, as new oligodendrocytes continue to be generated in the adult CNS, each of which produces dozens of sheaths (Hughes et al., 2018; Kang, Fukaya, Yang, Rothstein, & Bergles, 2010; Xiao et al., 2016). This phenomenon was visible during *in vivo* imaging in *Mobp-EGFP* mice as new EGFP-expressing (EGFP+) oligodendrocytes appeared within the imaging field (Figure 1E, F, H), continuously adding to the baseline oligodendrocyte population (Figure 1H; Supplementary Video 2). When mice were fed cuprizone for three weeks, > 90% of the baseline population of oligodendrocytes within the upper layers of cortex degenerated (94.2 ± 0.05%; N = 6 mice, mean ± SEM) (Figure 1E,G,I; Supplementary Video 3), with most cell loss occurring after cessation of cuprizone exposure (Figure 1I,J). New oligodendrocytes initially appeared rapidly during the recovery phase (Figure 1K); however, this burst of oligodendrogenesis was not sustained, and as a consequence, only about half of the oligodendrocytes (55.2 ± 0.03%) were replaced after nine weeks of recovery. Extrapolating from the rate of addition from the last recovery time-point (3.5 ± 0.5% per week, Figure 1K), it would take approximately three additional months (∼12.8 weeks) to achieve the density of oligodendrocytes prior to cuprizone, which is ∼two-fold greater in middle-aged cortices (Hughes et al., 2018). Thus, these young mice would be > 13 months old by the time they achieved full recovery, assuming a constant rate of addition. However, as oligodendrogenesis declines with age in both control and cuprizone-treated mice (Figure 1K) relative to age-matched control mice, cuprizone-treated mice may never reach a normal oligodendrocyte density after demyelination. These results indicate that although a prominent regenerative response is initiated in cortical gray matter, oligodendrocyte regeneration is much slower (Baxi et al., 2017) and less complete than in white matter (Baxi et al., 2017; Gudi et al., 2009; Jeffery & Blakemore, 1995; Matsushima & Morell, 2001).

### Layer specific differences in cortical remyelination

The cerebral cortex is a highly layered structure, in which genetically and morphologically distinct neurons form specialized subnetworks (Lodato & Arlotta, 2015), raising the possibility that they may adopt different myelination patterns to optimize circuit capabilities (Micheva et al., 2016; Stedehouder et al., 2017; Tomassy et al., 2014). Indeed, myelination patterns are highly non-uniform across the cortex (Supplementary Figure 2) and both oligodendrocyte density and myelin content increase with depth from the cortical surface (Hughes et al., 2018) (Figure 1C). However, it is not known whether the capacity for myelin repair is comparable between cortical layers. To examine depth-dependent changes in oligodendrogenesis, we subdivided imaging volumes into three 100 μm zones starting from the pial surface (Figure 2A-C). In control mice, the proportional rates of oligodendrocyte addition (top: 2.62 ± 0.37%; bottom: 2.93 ± 0.26% per week) were similar between the top zone (0-100 μm) and bottom zone (200-300 μm), despite their dramatically different oligodendrocyte densities (0-100 μm: 17 ± 2 cells; 200-300 μm: 79 ± 10 cells, N = 11 mice @ baseline) (Figure 2D,E); the only exception was for times when no cells were incorporated into the top zone (Figure 2E, *p =* 0.0036 @ 5 weeks, N-way ANOVA with Bonferroni correction for multiple comparisons). These findings suggest that oligodendrocyte enrichment in deeper layers occurs early in development, but then proceeds at similar rates across cortical layers in adulthood.

**Figure 2:**
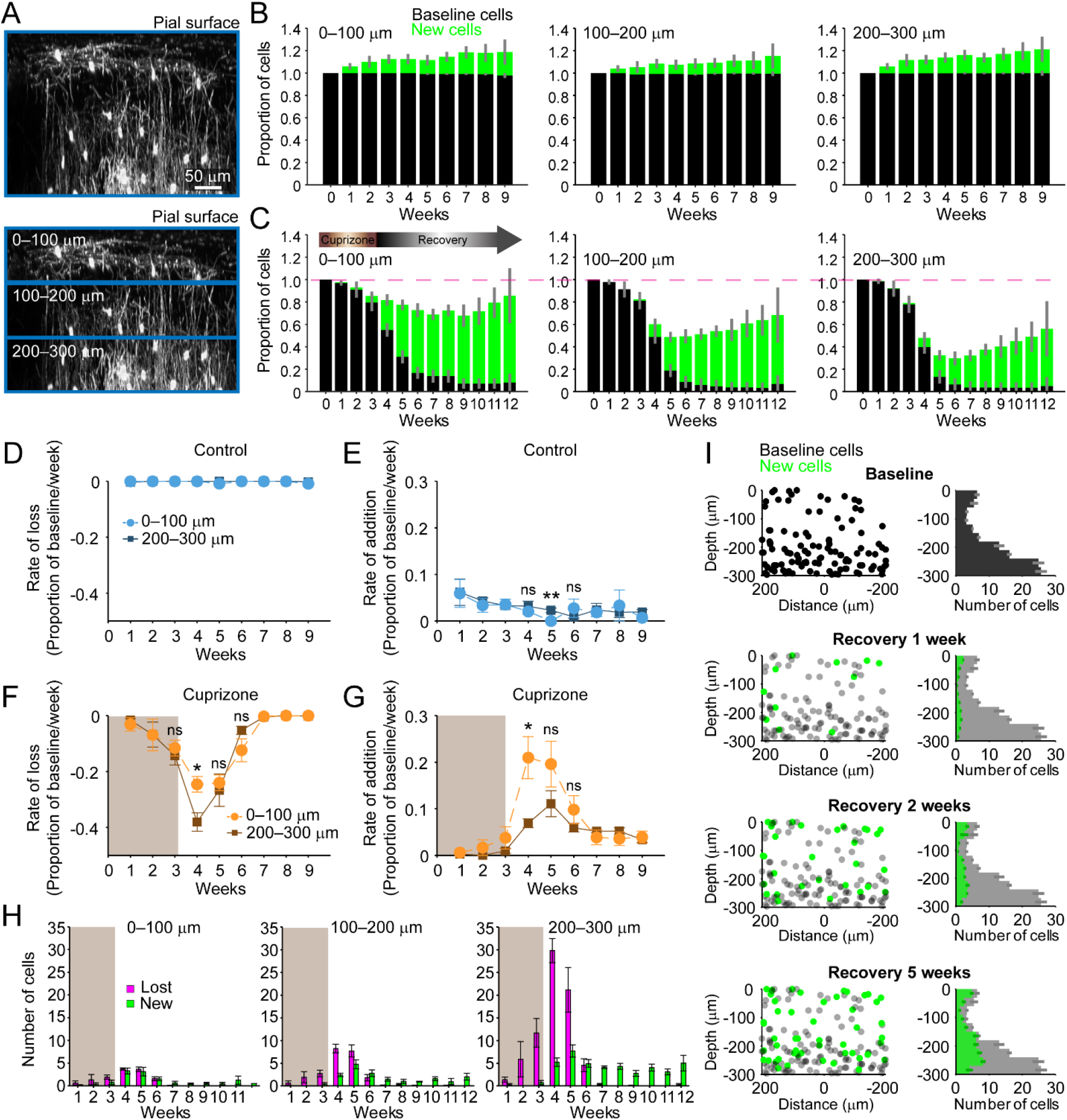
Regeneration of oligodendrocytes is incomplete in lower cortical regions. A: The fate of individual oligodendrocytes were determined within the same cortical volume (425 μm × 425 μm × 300 μm) over time that was divided into 0-100, 100-200, or 200-300 μm zones, (example maximum intensity Y projection is 425 μm × 150 μm × 300 μm). B-C: Mice were given either control diet (B) or cuprizone-supplemented diet (C), and individual cells present at baseline (black) or newly appearing over the time-course (green) were tracked longitudinally. The average cell counts per volume are depicted as a proportion of baseline number of cells (B: N = 5 control mice; C: N = 6 cuprizone-treated mice; with the same number of mice imaged at each time point as depicted in Fig. 1H,I). D-G: The rate of cell loss (D, F) or addition (E,G) relative to the baseline population of oligodendrocytes in each zone are depicted for each imaging time-point as a function of cortical depth (0-100 v. 200-300 μm zones), over 9 weeks of imaging in control (D-E) or cuprizone-treated (F-G) mice. Treatment with cuprizone-supplemented chow denoted by shaded background (F-G). Cells are rarely lost in control regions (D), and the rate of cell addition in the 0-100 (light blue circles) v. 200-300 μm (dark blue squares) zones are similar (except @ week 5, *p* = 0.0036). In the bottom 200-300 μm zone (F-G, in cuprizone-treated mice (brown squares), the rate of oligodendrocyte loss (F) is significantly greater at week 4 relative to the top (0-100 μm, orange circles) zone (*p* = 0.0443), and the rate of addition is significantly lower (G; *p* = 0.0364). H: The mean number of oligodendrocytes lost (magenta) and added (green) at each imaging time-point, comparing the 0-100 μm and 200-300 μm zones. I: The distribution of baseline (gray) and new (green) oligodendrocyte cell bodies relative to the center of an example region volume (left-side panels; 425 μm × 425 μm × 300 μm) and for all regions quantified (right-side panels; mean values from N = 6 mice) relative to the pial surface (0 on y-axis) is shown at baseline and at recovery weeks 1, 2 and 5 (corresponding to week 4, 5, and 8 of imaging weeks from A-H). Mean values depicted with error bars as standard error of the mean, and statistical tests are N-way ANOVA with Bonferroni correction for multiple comparisons.

In an efficient regenerative process, cell generation would be matched to cell loss; however, we found that oligodendrocyte regeneration was not proportional to their original density. Oligodendrocyte recovery was highly efficient in the top 100 μm zone, reaching 85.2 ± 0.17% of the baseline oligodendrocyte population nine weeks after cuprizone, but only 55.5 ± 0.11% of baseline after nine weeks in the bottom 100 μm zone (Figure 2C). The peak of this oligodendrocyte loss and replacement occurred during the first two weeks of recovery post-cuprizone (Figure 1J,K, Figure 2F,G). During this period, there was both a proportionally higher rate of cell loss and lower rate of cell replacement in the bottom 100 μm zone compared to the upper zone (Figure 2F,G) (loss @ week 4, *p* = 0.044; addition @ week 4, *p* = 0.036, N-way ANOVA with Bonferroni correction for multiple comparisons), indicating that the ability to balance cell loss with replacement declines with depth during this initial period. Although proportionally lower, more oligodendrocytes were generated per week in deeper layers (Figure 2H, green bars) (0-100 μm: 0.5-3.3 cells/week; 200-300 μm: 2.8 – 7.7 cells/week, *p* = 5.82 × 10^−4^ @ week 4, *p* = 0.046 @ week 5, N-way ANOVA with Bonferroni correction for multiple comparisons), but this enhanced oligodendrogenesis was not sufficient to compensate for the greater cell loss (Figure 2H, magenta bars). This relative lag in regeneration is clearly visible in maps indicating where newly generated oligodendrocytes were formed with regard to depth and their corresponding density histograms (Figure 2I, green circles and bars). This analysis highlights that the absolute number of oligodendrocytes integrated was remarkably similar in the top and bottom zones during the first few weeks of recovery, suggesting that there are factors that suppress regeneration of needed oligodendrocytes in deeper cortical layers.

A decrease in the pool of progenitors or formation of reactive astrocytes are potential candidates to impair oligodendrocyte regeneration in deeper cortical layers. Although OPCs are slightly less abundant in the deeper layers compared to the surface 100 μm (Supplementary Figure 3A,C,D) (*p* = 2.084 × 10^−13^, N-way ANOVA, by depth), there was no evidence of persistent OPC depletion during recovery (Supplementary Figure 3D,F) (*p* = 0.086, Kruskal-Wallis one-way ANOVA). Reactive astrocytes, a known pathological feature of both the cuprizone model and cortical demyelinating lesions (Chang et al., 2012; Skripuletz et al., 2008), can impair OPC differentiation by secreting cytokines (Kirby et al., 2019; Su et al., 2011; Zhang et al., 2010), but whether reactive astrocytes impair recovery differently as a function of cortical depth is unknown. GFAP+ astrocytes are relatively rare in the deeper (200-500 μm) regions of cortex in naïve mice; however, following cuprizone-treatment, their number increased nearly ten-fold (*p* = 5.86 × 10^−16^, N-way ANOVA, by time-point). This enhanced GFAP expression remained elevated throughout the recovery period (Supplementary Figure 3G) (p = 0.006, Kruskal-Wallis one-way ANOVA, with Fisher’s correction for multiple comparisons) and they retained a reactive morphology, even after five weeks of recovery (Supplementary Figure 3A,B,E,G). In contrast, astrocytes in the surface 100 μm, while exhibiting higher GFAP immunoreactivity at baseline, exhibited only a transient increase that was not sustained past two weeks of recovery (Supplementary Figure 3E). These findings highlight that the gliotic response to demyelination varies in magnitude and duration across the cortex, which may impair the recovery of gray matter regions with higher myelin content.

### Regeneration alters the pattern of myelination in the cortex

Myelination patterns are distinct among different neuron classes in the cortex (Micheva et al., 2016; Stedehouder et al., 2017; Tomassy et al., 2014), and can be discontinuous, in which myelin segments along individual axons are separated by long regions of bare axon (Tomassy et al., 2014). Even these isolated myelin segments are highly stable (Hill et al., 2018; Hughes et al., 2018), suggesting that preservation of these patterns is important to support cortical function, and therefore that recreation of these patterns should be a goal of the regenerative process. We hypothesized that given the sparseness of myelination in the cortex, new oligodendrocytes generated after demyelination should be formed in locations very close to the original population. To explore the spatial aspects of oligodendrocyte replacement, we mapped the distribution of oligodendrocytes within 425 μm × 425 μm × 300 μm volumes in 3-D prior to and after recovery from cuprizone, as well as the distribution of new cells generated in control mice (Figure 3A,B, green circles). Unexpectedly, these maps revealed that the cell bodies of regenerated oligodendrocytes occupied positions distinct from the original cells (see also Figure 1E). To quantify displacement of newly formed oligodendrocytes from original locations, we determined the Euclidean distance between nearest neighbors (nearest neighbor distance, NND) of every oligodendrocyte in this region at baseline and at five weeks of recovery (or eight weeks in control). Small changes in cell position were observed for oligodendrocytes in control mice (Figure 3C, Stable cells); however, addition of new oligodendrocytes to the cortex over this period did not significantly alter NND (All cells vs. Stable cells, 0-300 μm: *p* = 0.284, N-way ANOVA with Bonferroni correction for multiple comparisons). In contrast, regenerated oligodendrocytes added to the cortex of cuprizone-treated mice were significantly displaced relative to original locations (Figure 3C, All Cells vs. Stable cells) (*p* = 8.95 × 10^−7^, N-way ANOVA with Bonferroni correction for multiple comparisons). This apparent rearrangement of oligodendrocyte position was not due to differences in image quality or registration across the time series, or to changes in the structure of the tissue due to cuprizone exposure, as oligodendrocytes that survived in cuprizone exhibited the same average displacement as cells in control mice over the course of eight weeks of imaging (Figure 3C, Stable cells, Control vs. Regenerated, *p* > 0.05, N-way ANOVA with Bonferroni correction for multiple comparisons). This increase in NND was also not due to incomplete recovery from demyelination, as NND in the upper zone (0-100 μm) was greater than the average NDD across the whole volume (0-300 μm), despite the proportionally greater regeneration in layer I (Figure 2C). These results demonstrate that regenerated oligodendrocytes occupy locations within the parenchyma that are distinct from those present at baseline, and therefore may establish a new pattern of myelination.

**Figure 3:**
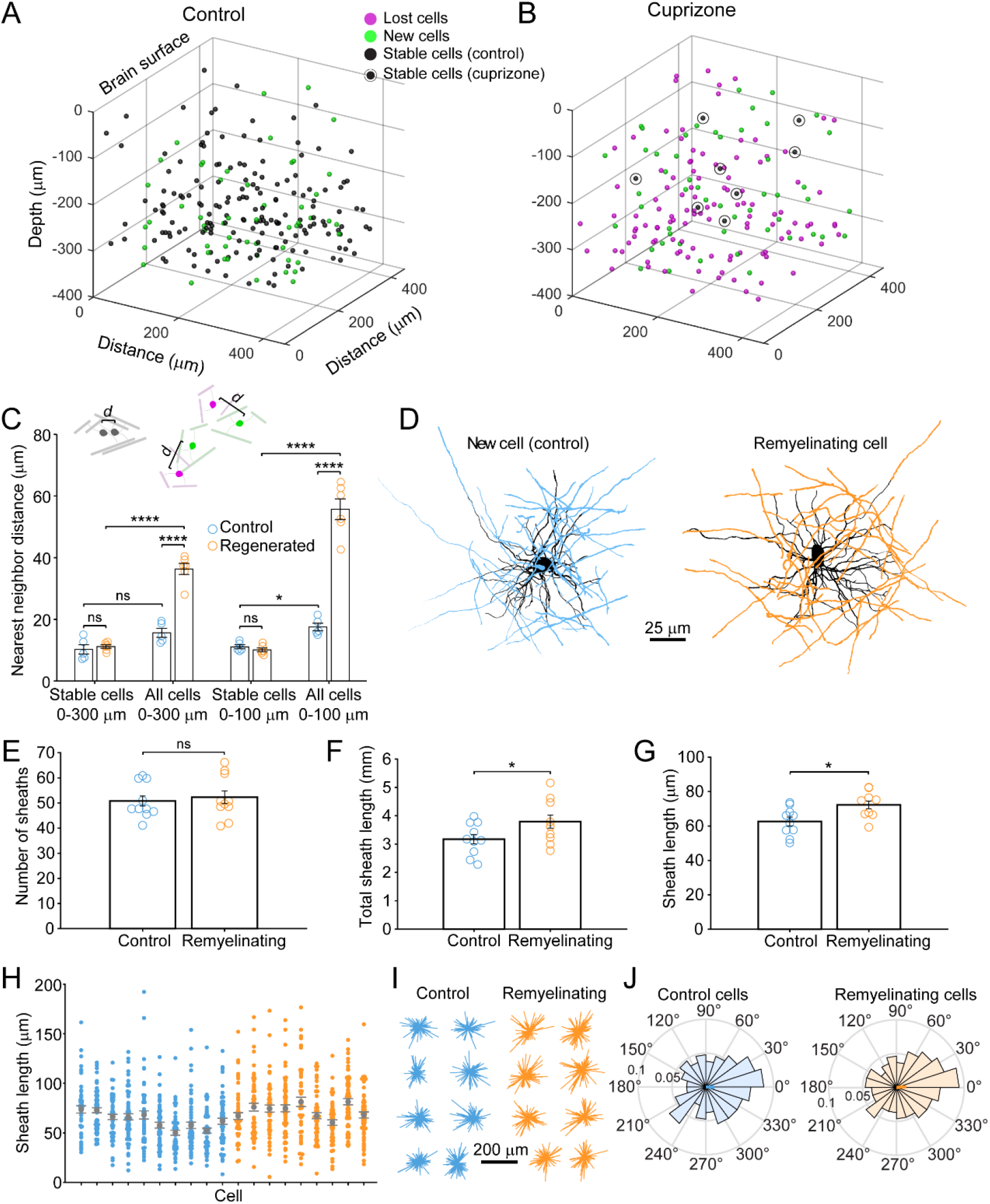
Remyelinating oligodendrocytes appear in novel locations but maintain morphology of oligodendrocytes generated in cuprizone-naïve mice. A-B: The location for all oligodendrocyte cell bodies present within the same cortical volume (425 μm × 425 μm × 300 μm) at baseline and after 8 weeks of two photon *in vivo* imaging are plotted and overlaid in 3-D. The fate of each cell over this time-course is designated as stable (black), lost (magenta), new (green). Example volumes for control (A) and cuprizone-treated (B) cortex are shown. C: The average 3-D nearest neighbor distance (NND) for lost, new, and stable cells, comparing baseline and last (8 weeks) imaging time-point locations, as plotted in A-B, were determined. Stable cells in both control (blue) v. cuprizone-treated (orange) cortices are minimally displaced (schematized by two gray oligodendrocytes displaced by *d* distance) in both the full 425 μm × 425 μm × 300 μm (0-300 μm; *p* = 4.64) volumes and the surface 425 μm × 425 μm × 100 μm (0-100 μm, *p* = 2.36) volume. New cells added into control cortex did not significantly alter NND in the full 0-300 μm region (controls cells (blue), all cells 0-300 μm v. stable cells 0-300 μm, *p* = 0.284), but did so in the top 0-100 μm zone, (controls cells (blue), all cells 0-100 μm v. stable cells 0-100 μm, *p* = 0.0137). The displacement of oligodendrocytes in cuprizone-treated cortex (schematized displacement (*d)* between lost (magenta) and new (green) cells) was greater in cuprizone-treated cortex (regenerated cells (orange), all cells 0-300 μm v. stable cells 0-300 μm; *p =* 8.95 × 10^−^*^7^*), despite near complete replacement of cells in the top 0-100 μm zone (regenerated cells (orange), all cells 0-100 μm v. stable cells 0-100 μm; *p =* 9.12 × 10^−7^). All comparisons in C are N-way ANOVA with Bonferroni correction for multiple comparisons. D: Every cell body (black), process (black) and myelin sheath (blue, control (10 cells, N=3 mice); orange, cuprizone (9 cells, N=3 mice)) belonging to newly appearing oligodendrocytes in cortical layer I (top zone) were traced using Simple Neurite tracer (Image J) on day of appearance. Shown are examples of maximum intensity projections of these rendered pseudocolored tracings. E: Newly formed control and remyelinating oligodendrocytes have similar numbers of myelin sheaths (*p* = 0.628, unpaired two-tailed t-test). F-G: The total length of myelin formed by a new cell (F) and the average length per sheath (G) for new control (blue) and remyelinating (orange) cells. Remyelinating cells overall make significantly more myelin (F; *p* = 0.041, unpaired two-tailed t-test) and individual myelin sheaths are longer (G; *p* = 0.012, unpaired two-tailed t-test). H: The distribution of sheath lengths per cell traced is plotted (each column represents the distribution of sheath lengths for an individual cell at 12-14 days from appearance (control, blue; remyelinating, orange, with mean and SEM in gray). I-J: Vectors were calculated from the cell body extending to each paranode of a reconstructed oligodendrocyte at day 12-14, then *x* and *y* vector components were summed and oriented to same direction of the vector sum. These are plotted (I) in 2-D for each cell traced (Control, N = 10 cells from 3 mice; Cuprizone, N = 9 cells from 3 mice), and were not significantly different from uniformly radial (J, Control, –0.047 ± 1.30 (std) rad, *p* = 0.400; Regenerated, 0.076 ± 1.30 (std) rad, *p* = 0.256, Hodges-Ajne test of non-uniformity). The average circular morphologies for new control or remyelinating cells (J, same cells as in I) exhibit similar degrees of circularity (p > 0.1, k = 462, Kuiper two-sample test).

The new locations of oligodendrocytes after a demyelinating event may not preclude these cells from myelinating the same axonal segments, as cortical oligodendrocytes can extend long cytoplasmic processes to form sheaths distant from the cell body (Chong et al., 2012; Murtie, Macklin, & Corfas, 2007). To determine whether regenerated oligodendrocytes restore the pattern of myelination by extending longer processes, we reconstructed their complete morphologies and compared these to oligodendrocytes generated at the same age in control mice. For new oligodendrocytes that appeared in layer I in control (10 cells, N = 3 mice) or cuprizone-treated mice (9 cells, N = 3 mice), each process extending from the cell body and every myelin sheath connected to these processes were traced when the cells first appeared in the imaging volume (Figure 3D), and for every 2 - 3 days for up to 14 days. In both control and cuprizone treated mice, newly generated oligodendrocytes exhibited an initial period of refinement after first appearance (governed by the onset of *Mobp* promoter activity and EGFP accumulation in the cytoplasm), characterized by sheath extensions and retractions (Supplementary Fig. 4A-D), pruning of entire myelin sheaths and removal of cytoplasmic processes (Supplementary Fig. 4E-H), before reaching a stable morphology (Supplementary Fig. 4I,J). The final position of the cytoplasmic process along the length of the sheaths (the likely point of sheath initiation) was randomly distributed along the length of the internode (Supplementary Fig. 4K,L). Notably, the initial dynamics of these newly formed oligodendrocytes are remarkably similar to those described in the developing zebrafish spinal cord (Auer, Vagionitis, & Czopka, 2018; Czopka, Ffrench-Constant, & Lyons, 2013), suggesting that the developmental sequence of oligodendrocytes is both highly conserved and cell intrinsic.

This time-lapse analysis revealed that the morphological plasticity of newly formed oligodendrocytes lasts for more than one week in the cortex (Supplementary Figure 4I,J); therefore, comparisons between control and regenerated oligodendrocytes were made from cells 12-14 days after first appearance when they reached their mature form. Although regenerated oligodendrocytes had access to much more axonal territory, they produced similar numbers of sheaths (Figure 3E) (Control: 53 ± 3 sheaths, 10 cells, N = 3 mice; Regenerated: 51 ± 2 sheaths, 9 cells, N = 3 mice, *p* = 0.628, unpaired two-tailed t-test). However, regenerated cells formed more total myelin (Figure 3F) (Total sheath length: Control, 3.17 ± 0.16 mm; Regenerated: 3.80 ± 0.23 mm, *p* = 0.041, unpaired two-tailed t-test) by extending slightly longer sheaths (Figure 3G) (average sheath length: Control, 62.6 ± 2.6 μm; Regenerated, 72.3 ± 2.2 μm, *p* = 0.012, unpaired two-tailed t-test), despite having similar distributions of sheath lengths (Figure 3H).

If regenerated oligodendrocytes reach the same axonal regions from a greater distance away, their processes should be more polarized; however, 2-D maps of sheaths arising from single cells, revealed that they exhibited a similar radial morphology (Figure 3D). To obtain a quantitative measure of polarization, vectors were calculated from the cell body to each paranode (Figure 3I) and mean radial histograms for new and remyelinating cells were calculated (Figure 3J). The average extent of polarization for control and regenerated cells was not significantly different from uniformly radial (Control, –0.047 ± 1.30 (std) rad, *p* = 0.400; Regenerated, 0.076 ± 1.30 (std) rad, *p* = 0.256, Hodges-Ajne test of non-uniformity) and sheaths of new cells in control and those regenerated after cuprizone exhibited similar degrees of circularity (*p* > 0.1, k = 462, Kuiper two-sample test). Thus, regenerated oligodendrocytes formed in an environment with greater myelination targets have morphologies remarkably similar to cells added to existing myelinated networks in naïve mice, suggesting that oligodendrocyte structure is shaped by strong cell intrinsic mechanisms.

To estimate the extent of myelin sheath recovery in the somatosensory cortex, we measured the overlap in cell territory between baseline and regenerated cells. Territories of individual oligodendrocytes which existed at baseline, those generated in control, and those regenerated following cuprizone were estimated using an ellipsoid centered at the center of mass of the sheath arbor with the smallest radii in the *x-y* (oriented parallel to the pia) and *z* planes (oriented perpendicular to pia) that encompassed at least 80% of the total sheath length. These radii were averaged across all cells per condition to generate model ellipsoids (Figure 4A), representing the volume available to be myelinated by an individual oligodendrocyte. As expected from the slightly longer myelin sheaths produced by remyelinating cells, their average cell territory was significantly larger than control cells (Figure 4B) (*x-y*; Control: 75.7 ± 2.1 μm; Regenerated: 85.2 ± 2.8 μm, *p* = 0.025, one-way ANOVA). Model ellipsoids were then centered on the cell body coordinates in layer I from control and cuprizone-treated mice (examples shown in Figure 4C,D) and their degree of territory overlap determined. This analysis revealed that regenerated oligodendrocytes in cuprizone exposed mice exhibit only 59.1 ± 5.8 % territory overlap (Range: 37.2 – 79.0%, N = 6) with oligodendrocytes that originally populated these regions of the cortex. This convergence was slightly, but not significantly, higher than predicted if the same number of regenerated cells were placed at random in the volume (44.3 ± 4.7%; *p* = 0.078 by t-test) (Figure 4E), suggesting that local factors may influence which progenitors differentiate. Moreover, because regenerated cell territories were larger (Figure 4B), they enclosed an average volume of 116 ± 14.1% of the total territory at baseline (Range: 87.4 – 181%) (Figure 4F). These data and those obtained from the NND analysis indicate that although regenerated oligodendrocytes tend to be formed close to the sites of original cells, they unexpectedly do not completely overlap with the baseline territory; thus, regenerated oligodendrocytes are unlikely to access to the same complement of axons to myelinate, and could potentially myelinate novel axon segments.

**Figure 4:**
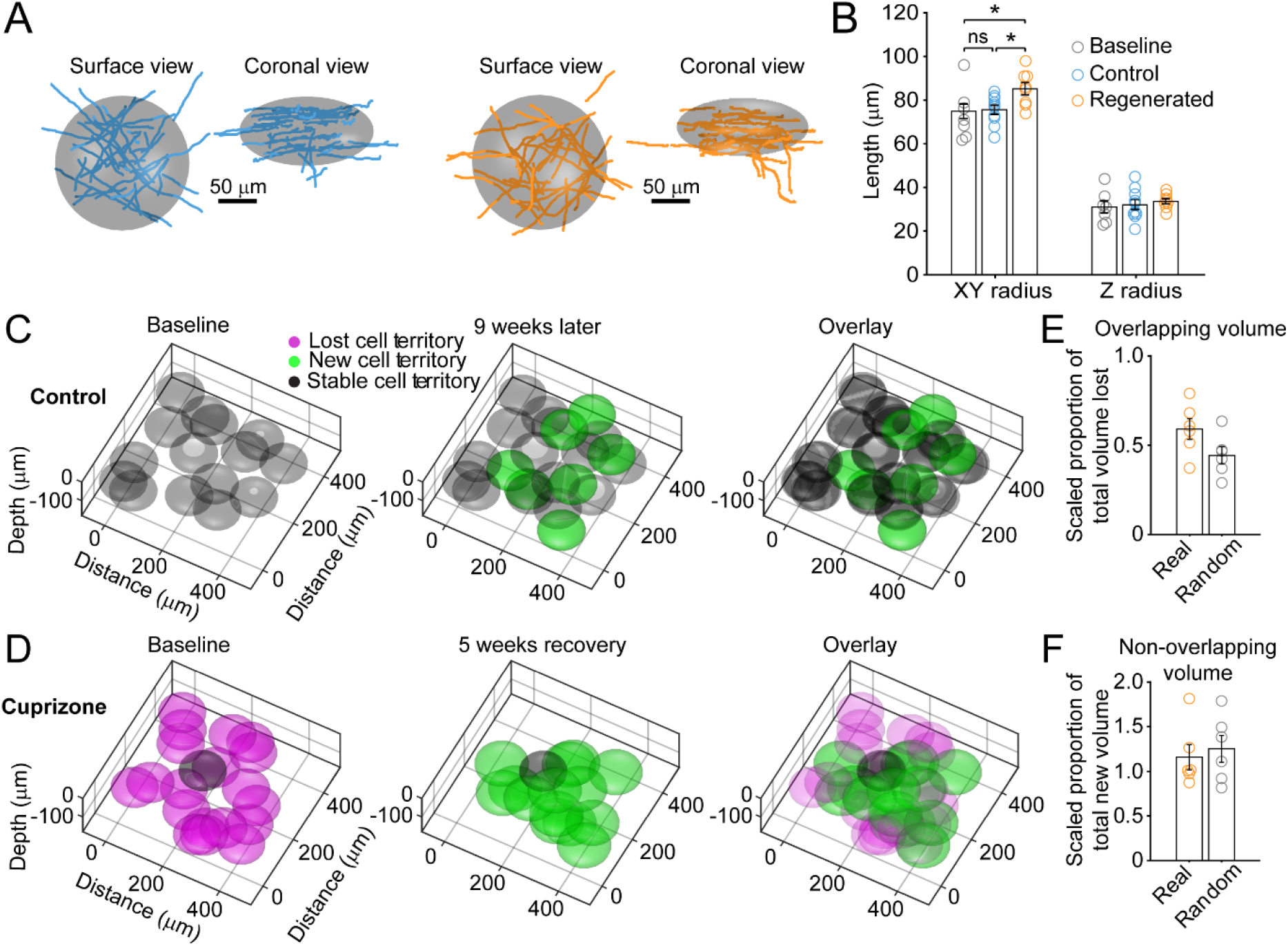
Individual remyelinating oligodendrocytes myelinate cortical territory that is distinct from lost oligodendrocytes. A-B: The best fit ellipsoid (E) that encompasses 80% of all myelin sheaths of an individual oligodendrocyte (examples shown in A for new control (blue) and remyelinating (orange) cells in the surface 0 -100 μm zone (for *x-y* radius) and coronal (for *z* radius) views) was calculated and the average value ((Control (blue), N = 10 cells from 3 mice; Cuprizone (orange), N = 9 cells from 3 mice)) is plotted in B ((gray, cells at baseline (N = 7 cells from 4 mice)). Circles represent individual cells). Remyelinating cells were significantly wider (*x-y* radius) than new control cells (*p* = 0.025, one-way ANOVA). C-D: The average best-fit ellipsoids for control (C) or remyelinating cells (D) as calculated in (B) were plotted in example regions from the top 0-100 μm of cortex (425 μm × 425 μm × 100 μm) based on location of cell bodies. E-F: The proportion of total territory volume at baseline that was overlapped by regenerated oligodendrocytes (scaled to take into account differences in numbers of baseline and regenerated cells; see Methods), and the additional volume encompassed by regenerated oligodendrocyte territory relative to total baseline volume (F), for 0-100 μm regions in cuprizone-treated (orange, N = 6 mice) and compared to the volumes predicted if the same number of regenerated cells appeared at random within the same volumes (gray). Regenerated oligodendrocyte territories partially overlap with baseline volume (E; 59.1%, *p* = 0.0778 by one-way ANOVA) and encompass novel territory (F; 115%, *p* = 0.662 by one-way ANOVA), at a similar proportion to regenerated cells placed at random.

### Regeneration of specific myelin segments

Although oligodendrocytes are formed in new locations during recovery from demyelination, they have the opportunity to regenerate specific myelin sheaths in areas where they extend processes into previously myelinated territories. To assess the extent of sheath replacement in the somatosensory cortex, we acquired high resolution images in layer I in *Mobp-EGFP* mice, allowing assessment of the position and length of individual myelin internodes. We randomly selected 100 μm × 100 μm × 100 μm volumes in the image stacks (425 μm × 425 μm × 100 μm), and traced the full length of internodes that entered the volume at baseline and at the five-week recovery time point in control (N = 5) and cuprizone-treated mice (N = 5) (Figure 5A). Internodes were classified as *stable* (supplied by the same cell at baseline and recovery), *lost* (present at baseline and absent in recovery), *replaced* (present at baseline, lost and then replaced by a new cell), or *novel* (not present at baseline, but present in recovery). Consistent with previous findings (Hill et al., 2018; Hughes et al., 2018), in control mice almost all internodes (99.1 ± 0.5 %) present at baseline remained after five weeks (Figure 5A, gray processes), demonstrating the high stability of myelin sheaths within the cortex. Generation of new oligodendrocytes within this area led to the appearance of a substantial proportion (25.8 ± 8.7 %) of novel sheaths (Figure 5A, green processes). In mice treated with cuprizone, 84.4 ± 2.7% of myelin sheaths were destroyed in this region (Figure 5B), but sheath numbers were restored to baseline levels after five weeks (Figure 5C). However, despite this apparent recovery of overall myelin content, only 50.5 ± 2.9% of specific internodes were replaced, 32.4 ± 1.2% were lost (not replaced) and 47.6 ± 9.9 % novel internodes were formed (N = 5 mice) (Figure 5D), indicating that regeneration leads to a dramatic change in the overall pattern of myelin within cortical circuits.

**Figure 5:**
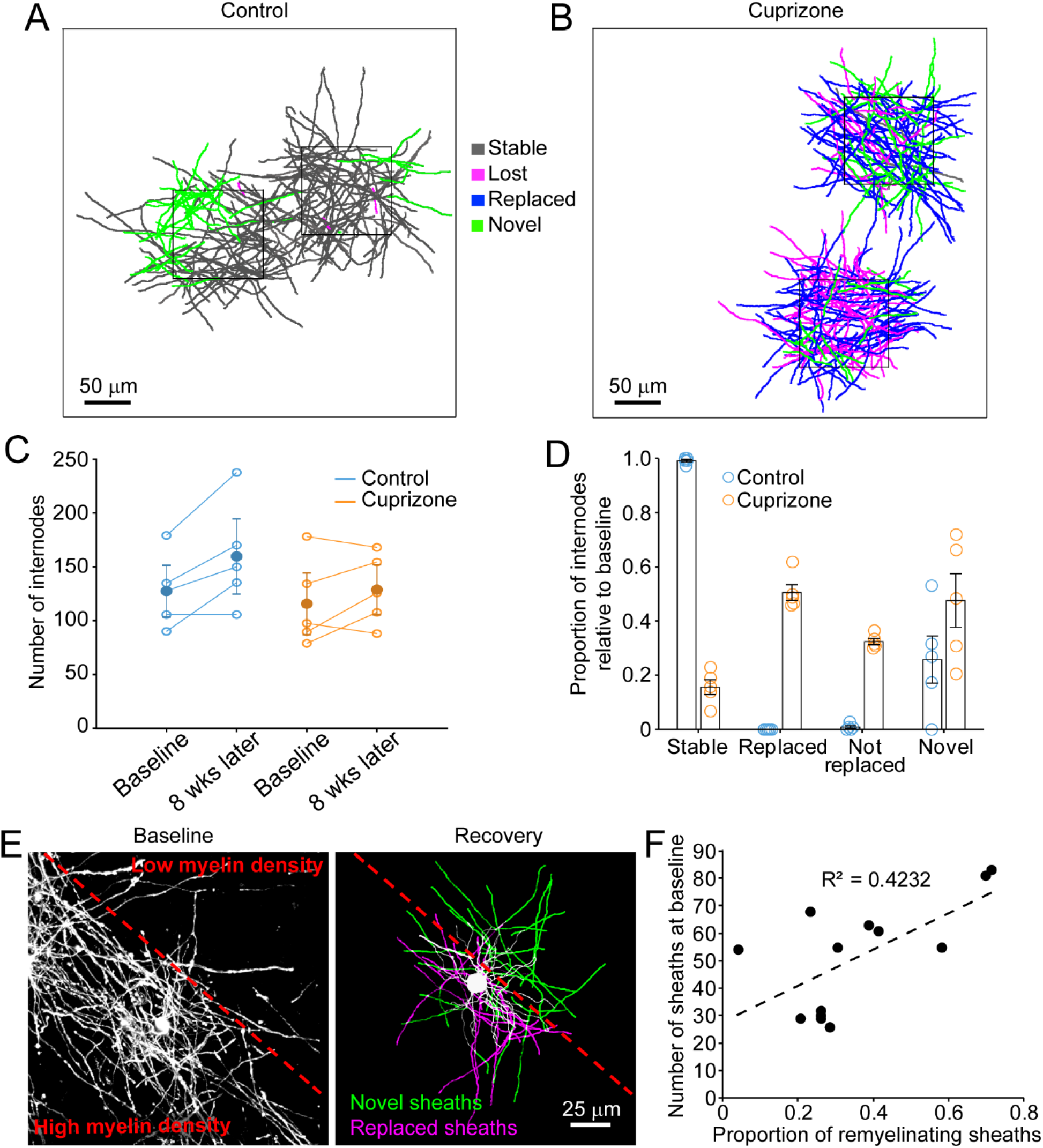
After oligodendrocyte loss, regeneration results in a new pattern of cortical myelin. A-B. Individual myelin sheaths which passed through a 100 μm × 100 μm × 100 μm volume within the top 0-100 μm zone were traced, and the fate of each sheath within the volume was determined to be stable (black), lost (magenta), replaced (blue) or novel (green) across the time series. Examples of traced myelin sheaths with pseudocolor designation from control (A) and cuprizone-treated (B) cortex are depicted. C: Total number of traced internodes at baseline and after 8 weeks of imaging are plotted (circles represent means for individual mice, with line connecting two time-points; mean for all mice is filled circle with SEM) for control (blue, N = 5), and cuprizone-treated (orange, N = 5) mice. D: Quantification of internode fates from cuprizone-treated (orange, N = 5) and control (blue, N = 5) mice (circles represent mean proportional values relative to baseline from individual mice). Error bars are standard error of the mean. E: Baseline panel on left is a maximum intensity projection (226 μm × 226 μm × 60 μm) illustrating the myelin sheaths at baseline, with a red dashed line demarcating an area of higher (lower left) v. lower myelin density (top right). Right panel is a rendering of a completely traced and pseudocolored new oligodendrocyte that appears in the region at 3 days recovery and forms myelin sheaths that either replace those lost (magenta) or are novel (green). F: The proportion of replaced sheaths per new oligodendrocyte were plotted as a function of the number of baseline myelin sheaths present within the territory (average remyelinating ellipsoid, see Figure 4A,B) of the new cells (13 remyelinating oligodendrocytes from N = 4 mice), correlation coefficient R^2^ = 0.4232.

Analysis of single oligodendrocytes revealed that the degree of sheath replacement varied considerably between regions. Oligodendrocytes regenerated in regions that originally had a high density of sheaths devoted a larger proportion of their sheaths to replacement (R^2^ = 0.4232, 13 cells, N = 4 mice) (Figure 5E,F). This phenomenon was most evident in cells that traversed boundaries of low and high myelin density. Nevertheless, rather than extending all processes into the previously highly myelinated area, they retained a radial morphology (Fig. 3I); processes that extended into densely myelinated areas often formed sheaths on previously myelinated axons, while processes that extended into the sparsely myelinated area typically formed novel sheaths (Figure 5E,F). These results suggest that remyelination may be opportunistic and based, at least initially, on chance interactions between processes and local axon segments. This finding is consistent with the highly radial morphology of premyelinating oligodendrocytes (Trapp, Nishiyama, Cheng, & Macklin, 1997), which would be optimized for local search of the surrounding volume rather than directed, regional targeting of subsets of axons.

Myelination patterns along axons in the cortex are highly variable, ranging from continuous to sparsely myelinated in the same region (Hill et al., 2018; Hughes et al., 2018; Micheva et al., 2016; Tomassy et al., 2014). To determine if there is a preference for replacing specific types of sheaths during recovery from demyelination, we classified sheaths present at baseline according to their neighbors: 0 neighbors within 5 μm (*isolated)*, one node of Ranvier (NOR) and one hemi-node (*1 neighbor*), or two NORs (*2 neighbors*) (Figure 6A,B; Supplementary Videos 4,5). In control mice this pattern was highly stable, with similar proportions of isolated and neighbored sheaths at baseline and eight weeks later (Figure 6C), a pattern that we showed previously is stable through middle-age (Hughes et al., 2018). Indeed, novel sheaths generated by new oligodendrocytes made roughly equal proportions of isolated and neighbored sheaths (Figure 6D), indicating that new sheaths were added to previously-unmyelinated axons, as well as next to existing sheaths on myelinated axons, visible as higher values in the upper right quadrant (relative to lower left quadrant) of the myelination matrix (Figure 6E). We then assessed the extent of sheath replacement five weeks after the end of cuprizone. Similar to that observed in control, oligodendrocytes in cuprizone-treated mice had similar proportions of isolated and neighbored sheaths as they did at baseline (Figure 6F). While lost (not replaced) and novel internodes exhibited both isolated and neighbored sheaths with equal proportion, the majority of replaced internodes had at least one neighbor (79.5 ± 2.5 %) (Figure 6G), visible in the bias towards higher values in the lower right quadrant in the myelination matrix (Figure 6H). These data suggest that regenerated oligodendrocytes preferentially formed sheaths on axons that were previously more heavily myelinated, suggesting that the factors which promote greater myelination of specific axons is preserved after demyelination.

**Figure 6:**
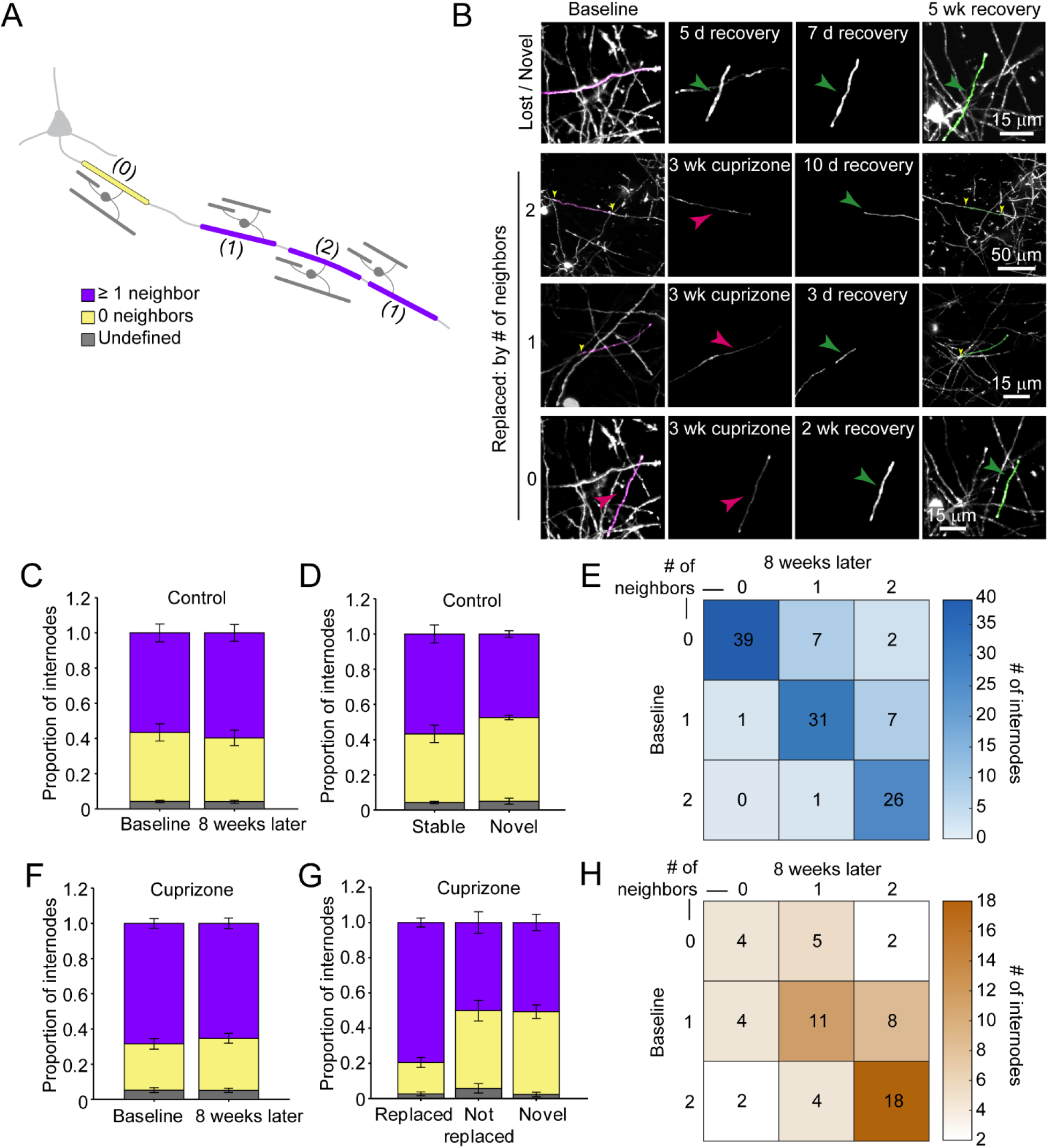
Myelin sheaths are more likely to be replaced if they had neighboring sheaths before demyelination. A: Schematic of intermittent myelination of cortical axons, designating each internode by number of flanking myelin sheath neighbors (0, yellow; 1 or 2, lavender; or undefined, gray). B: Example maximum intensity projections of lost (magenta) and new (green) myelin sheaths from cuprizone-treated mice. Top row is an example of a lost, but not replaced, sheath, as well as a novel sheath not present at baseline (green arrowhead). Remaining rows are examples of lost (magenta sheaths and arrowheads) and replaced internodes (green sheaths and arrowheads) that had 0, 1, or 2 neighboring sheaths at baseline. C, F: Comparison of mean proportion of internodes with at least one neighbor (lavender), isolated (yellow), or undefined (gray) within a 100 μm × 100 μm × 100 μm volume at baseline and 8 weeks later, from control (C, N = 5) and cuprizone-treated (F, N = 5) mice. Volumes from both control and cuprizone-treated conditions have the same relative proportion of isolated vs. ≥ 1 neighboring internode at both time-points (*p* > 0.05, N-way ANOVA with Bonferroni correction for multiple comparisons). D, G: Comparison of the mean proportion of internodes with 0 or ≥ 1 neighbor that are stable or novel (control, D) or replaced, not replaced, or novel (cuprizone, G). There is no significant difference in the proportion of isolated vs. ≥ 1 neighbor population between stable and novel sheaths in control (D, p > 0.05, unpaired two-tailed t-test with Bonferroni correction for multiple comparisons), nor between lost and novel sheaths in cuprizone (G, p > 0.05, unpaired two-tailed t-test with Bonferroni correction for multiple comparisons), but relatively more internodes with ≥ 1 neighbor were replaced in cuprizone-treated cortex (G, p = 5.93 × 10^−7^, unpaired two-tailed t-test with Bonferroni correction for multiple comparisons). E,H: The average number of internodes categorized by number of neighbors at baseline and then at final imaging time-point, plotted in a “myelination matrix” for control (E), where, there is a trend to increase number of neighbors over 8 weeks. In cuprizone-treated mice, the average number of neighbors for lost and replaced internodes are categorized by number of neighbors at baseline (lost internode) and after 5 weeks of recovery (8 weeks imaging, replaced internode). Internodes with more neighbors at baseline are more likely be replaced (largest average # of internodes in bottom right of matrix).

Analysis of individual sheath position revealed the remarkable spatial specificity with which sheaths were replaced. For both isolated sheaths and those along continuously myelinated segments, sheaths were often formed close to the same position along axons, resulting in NOR or hemi nodes at similar positions (Figure 7A, Supplementary Video 6). These results suggest that there may be persistent landmarks along axons indicating the prior position of nodes after myelin is removed (Figure 7B). To assess whether nodal components along cortical axons remain after demyelination, we performed post-hoc immunostaining on tissue from mice exposed to cuprizone for six weeks (longer than three weeks used above, to increase the length of time that axons were devoid of myelin) with antibodies against Caspr, βIV-spectrin, and Ankyrin-G, which together label paranodal and nodal regions along myelinated axons (Susuki, Otani, & Rasband, 2016). High resolution *z*-stacks (135 μm × 135 μm × 30 μm, 2048 × 2048 pixels) of layer V-VI were acquired in order to capture a large population of nodes (Figure 7C,D), as the upper layers of cuprizone-treated mice had inadequate numbers for quantification at this magnification. The full images were transformed into surfaces to allow automated quantification (see Methods) (Figure 7E,F). βIV-spectrin puncta were classified as “nodes” if they were flanked by Caspr+ puncta, “heminodes” if they were bound by only one side by Caspr, and “isolated” if no flanking Caspr was visible (Figure 7B). As expected, the distribution of nodes among these categories remained stable in control mice over two weeks (Figure 7G). In contrast, the characteristics of nodes changed dramatically after exposure to cuprizone, with a loss of nodes visible at four weeks, a time when ∼50% of oligodendrocytes had degenerated (Figures 1I, 7G). After six weeks in cuprizone, few Caspr+ puncta remained, and correspondingly there were few nodes or heminodes (Figure 7F,G). However, even at this later time point, many βIV-spectrin+ puncta were visible, with the greatest proportion of these isolated from Caspr, suggesting that βIV-spectrin remains clustered for a prolonged period, consistent with the hypothesis that many represent previous nodes (i.e. previously flanked by Caspr). Thus, although prior studies suggest that NOR components are rapidly disassembled and redistributed after demyelination (Coman et al., 2006; Craner et al., 2004; Dupree et al., 2004), these results indicate that βIV-spectrin, which links the underlying actin cytoskeleton to integral membrane proteins within the node/paranode, remains clustered along axons, which may provide the means to confine sheath extension to previously myelinated regions and allow precise restoration of myelin sheath position during regeneration.

**Figure 7:**
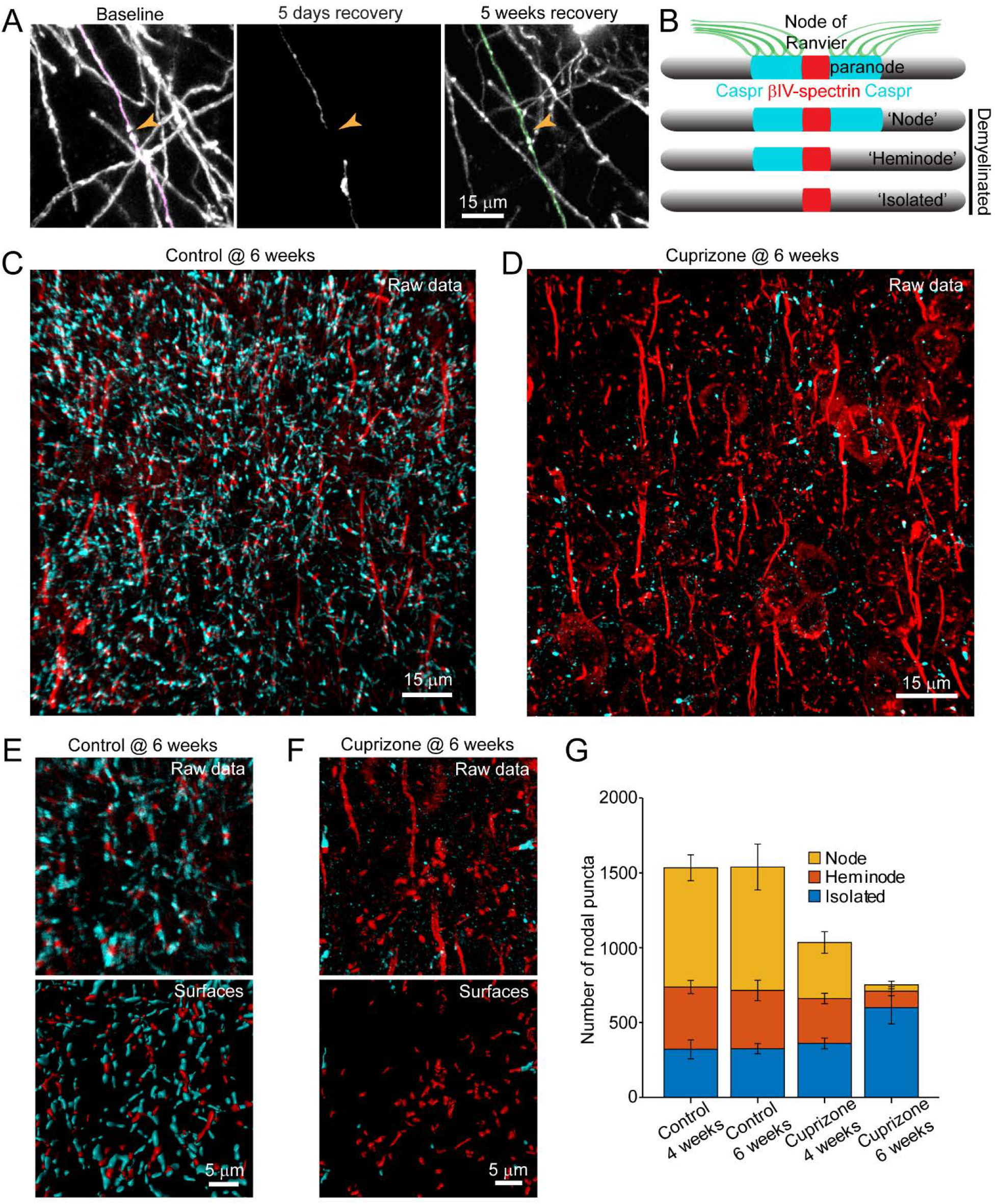
Structural components of the node of Ranvier persist after demyelination. A: Adult *Mobp-EGFP* mice were fed cuprizone-supplemented diet for three weeks and individual myelin sheaths were imaged and traced within a 100 μm × 100 μm × 100 μm volume to determine their fate. An example of myelin sheaths that were lost (magenta, baseline), and regenerated (green, 5 weeks recovery) are overlaid on maximum intensity projections from a longitudinally imaged somatosensory cortical region. These traced myelin sheaths are shown degenerating at 5 days recovery. Orange arrowhead denotes location of node of Ranvier at baseline and a similar position after 2 neighboring myelin sheaths are regenerated at 5 weeks of recovery. B. Schematic depicting axonal regions of a myelinated axon where βIV-spectrin (node of Ranvier, red) and Caspr (paranode, cyan) localize. After demyelination, puncta of βIV-spectrin can be found with 2 flanking Caspr puncta (“Nodes”), 1 flanking Caspr punctum (“Heminode”), or no nearby Caspr puncta (“Isolated”). C-D: Adult *Mobp-EGFP* mice were fed cuprizone-supplemented diet (D) or sham chow (C) for 6 weeks to induce loss of oligodendrocytes and inhibit formation of new oligodendrocytes. Coronal sections from brains of mice euthanized after 4 or 6 weeks of treatment were immunostained for βIV-spectrin and Caspr. E-F: Magnified views of immunostained somatosensory cortex (Raw data) from control (E) and cuprizone-treated (F) brains. Example post-processed “Surfaces” that were used to calculate nearest neighbor distances between βIV-spectrin puncta and Caspr puncta within 3.5 μm are shown for the same magnified images in E and F. Axon initial segments were excluded from surface rendering. G: Total number of βIV-spectrin puncta categorized as either Node, Heminode or Isolated. There are fewer overall βIV -spectrin puncta after 6 weeks of cuprizone (compared to controls (@4 or 6 weeks) or 4 weeks of cuprizone, but relatively more isolated puncta still present after 6 weeks of cuprizone treatment. (control @ 4 weeks, N = 3 mice; control @ 6 weeks, N = 3 mice; cuprizone @ 4 weeks, N = 4 mice; cuprizone @ 6 weeks, N = 4 mice). Bars are SEM.

## DISCUSSION

The organization of myelin in the cerebral cortex is remarkably diverse, with densities of myelin sheaths varying between cortical regions, between axons from different classes of neurons and even along individual axons within a given area. These patterns are established progressively over many months, creating an extended developmental time course that results in pronounced increases in myelin through adolescence and adulthood. Although progressive, oligodendrogenesis and myelination can be enhanced by life experience and may be critical to certain forms of learning (Gibson et al., 2014; Hughes et al., 2018; McKenzie et al., 2014). However, once formed, oligodendrocytes and their complement of myelin sheaths are extraordinarily stable, with cell survival and sheath position varying little over months and are resistant to environmental changes such as sensory enrichment, in accordance with the high stability of myelin proteins and the persistence of oligodendrocytes in the human CNS (Yeung et al., 2014). Experience dependent changes in myelin appear to occur primarily through addition of new sheaths arising from oligodendrogenesis (Hughes et al., 2018), but see also (Dutta et al., 2018; Gibson et al., 2014). The high stability of oligodendrocytes and their preferential myelination of specific neurons, such as parvalbumin interneurons in layers II/III (Micheva et al., 2016; Stedehouder et al., 2017), suggest that preserving the overall pattern of myelin is important to optimize and sustain the processing capabilities of these circuits. Consistent with this hypothesis, demyelination within the cortex in diseases such as MS is closely associated with cognitive impairment and increased morbidity. Moreover, many neurodegenerative diseases and affective disorders are associated with profound alterations in myelin (Gouw et al., 2008; Ihara et al., 2010; Stedehouder & Kushner, 2017), and recent studies indicate that demyelination triggers a dramatic decrease in excitatory synapses (Araújo et al., 2017; Werneburg et al., 2020), suggesting that disruptions in these sheaths may also influence both structural and functional aspects of circuits. Although preserving cortical myelination appears paramount, the highly variable patterns of myelin within cortical circuits and the sparseness of these neuron-glial associations create considerable challenges for regenerating specific myelin sheaths after injury or disease.

Here, we used two photon *in vivo* imaging to examine the destruction and regeneration of oligodendrocytes and myelin sheaths in cortical circuits of adult mice. These longitudinal, high resolution studies revealed features of the regenerative process in the cortex that were not previously known. First, oligodendrocytes were regenerated in new locations, yet had similar morphologies. Second, regenerated oligodendrocytes often formed sheaths along portions of axons that were previously unmyelinated, establishing a new pattern of myelination. Third, oligodendrocyte regeneration was not uniform across the cortex and became less efficient with depth from the cortical surface, in concert with the increasing density of myelin (prior to oligodendrocyte destruction) and enhanced gliosis. Fourth, in areas of territory overlap, regenerated oligodendrocytes were able to establish sheaths at similar positions along previously myelinated axons, indicating that positional cues persist along axons long after demyelination. Together, these findings reveal unexpected aspects of cortical remyelination, raising new questions about the mechanisms that impair oligodendrocyte regeneration in deeper cortical layers, the mechanisms that enable oligodendrocytes to identify and replace individual myelin sheaths and the long-term consequences of circuit level changes in myelination patterns.

### Regenerative potential of cortical OPCs

In regenerative processes, cell loss and cell generation are typically closely coupled, helping to ensure efficient replacement without energetically costly production of excess cells and further tissue disruption (Biteau, Hochmuth, & Jasper, 2011). If this scenario were to prevail in the CNS, the generation of new oligodendrocytes should be proportional to those lost, with much higher oligodendrocyte production occurring in deeper layers. However, our results indicate that oligodendrocyte replacement was remarkably constant across layers for many weeks after a demyelinating event (Figure 2G-I), with regeneration in deeper layers lagging behind that predicted for one-to-one replacement. This phenomenon could be explained if local environmental factors in deeper layers suppress OPC differentiation. The larger number of oligodendrocytes that degenerate in this area may inhibit regeneration by creating more inflammation and myelin debris, factors known to suppress oligodendrogenesis. Consistent with this hypothesis, reactive gliosis was more prominent and more prolonged in deeper layers of cortex (Supplementary Figure 3A,B,E,G). Nevertheless, even after extended recovery (> 9 weeks), oligodendrocyte density remained much lower in these regions than present at baseline. This regional suppression of oligodendrogenesis is particularly apparent when assessing the rate of cell addition, as the initial burst in oligodendrogenesis that occurs just following cell loss is not sustained despite the remaining cell deficit (Figure 2G-I). We predict that it would take approximately three additional months for the oligodendrocyte population to be fully regenerated, but this is only possible if a higher rate of oligodendrogenesis (∼3.5%) than observed in age-matched control brains (1.7%) is maintained. However, it is not clear whether the endogenous pool of OPCs could maintain this higher rate of differentiation (and resulting homeostatic OPC turnover) required for such a prolonged period of replacement. Although recent studies indicate that GLI1-expressing glial progenitors positioned within germinal zones along the lateral ventricles are mobilized in response to cuprizone-induced demyelination, forming new OPCs that migrate and differentiate into oligodendrocytes with higher probability than resident OPCs (Samanta et al., 2015), our results indicate that this recruitment is not sufficient in the short term to overcome existing inhibitory barriers within cortical gray matter.

The inability to recover fully from a demyelinating event raises the possibility that inflammation persists long after the initial trauma, or that OPCs in these regions are permanently altered as a result of exposure to this environment, if only for a short time (Baxi et al., 2015; Kirby et al., 2019). Like other progenitor cells, OPCs exhibit a decline in regenerative potential with age and can undergo senescence, a process that may be accelerated by exposure to inflammatory cytokines (Kirby et al., 2019; Neumann et al., 2019; Nicaise et al., 2019). It is also possible that there is a restricted time period during which OPCs can detect and respond to myelin loss; if there are inherent limits on OPC mobilization, as suggested by the uniform behavior of OPCs across cortical layers, then the inability to match the demand for new cells early may lead to prolonged deficits. In this regard, evidence that certain aspects of the inflammatory response strongly promote OPC differentiation (Kotter, Li, Zhao, & Franklin, 2006; Miron et al., 2013; Ruckh et al., 2012) raises the possibility that there is a critical time window for optimal repair. A more detailed spatial and cell-type specific profiling of inflammatory changes in the cortex after oligodendrocyte death may help clarify the role of inflammatory changes in this impaired regeneration.

An unexpected feature of oligodendrocyte regeneration in the cortex was that new cells were not formed in the same locations as the prior oligodendrocytes, even though the high density, regular spacing and dynamic motility of OPCs seem ideally suited to optimize placement of new oligodendrocytes. Inhibitory factors generated as a consequence of oligodendrocyte degeneration, such as myelin debris (Kotter et al., 2006), axonally expressed factors (LINGO1) (Mi et al., 2005), extracellular matrix components (chondroitin sulfate proteoglycans) (Lau et al., 2012), and reactive astrocytes (Back et al., 2005) may create a local zone of exclusion at sites of cell death reducing the probability of OPC differentiation.

Complete reconstruction of individual oligodendrocytes revealed that regenerated cells underwent a similar period of structural refinement over a period of ∼ 10 days and ultimately formed a comparable number of sheaths (Figure 3D,E, Supplementary Figure 4A-J). Although oligodendrocytes have the capacity to form long processes, they did not always extend their cytoplasmic processes to reach sites of original myelination, instead creating sheaths within a local territory similar to that of normal oligodendrocytes (Figure 3D,I), with only a 15.5% increase in total sheath length (Figure 3F). As regenerated oligodendrocytes are formed in an environment with an apparent surplus of receptive axons, these findings suggest that the size and shape of oligodendrocytes is profoundly limited by cell intrinsic mechanisms.

### Reparative potential of surviving oligodendrocytes

Recent studies of postmortem human tissue from MS patients have raised the intriguing possibility that remyelination may occur through the reformation of myelin sheaths by oligodendrocytes that survive autoimmune attack, rather than from *de novo* oligodendrogenesis (Yeung et al., 2019). This hypothesis is based on evidence that “shadow plaques”, which are classically considered to represent partially remyelinated axons (Lassmann, Brück, Lucchinetti, & Rodriguez, 1997), did not appear to contain many newly born oligodendrocytes, as assessed using C^14^–based birth dating; the progressive decline in atmospheric C^14^ levels following the cessation of atomic testing in the 1950s allow the date of last cell division to be estimated the amount of C^14^ present. Although *in vivo* imaging studies indicate that oligodendrocytes can be generated through direct differentiation of OPCs without cell division (Hughes et al., 2013), potentially confounding measures of cell age based on proliferation, these results nevertheless posit that significant new myelin can be created by existing oligodendrocytes. In our studies, the few oligodendrocytes that survived cuprizone did not contribute substantial new myelin. However, the cuprizone model does not fully recapitulate the pathology of cortical lesions observed in MS. In particular, demyelination of the upper layers of the cortex in autopsy samples from MS patients are correlated with regions of leptomeningeal inflammation, composed of B and T cells that secrete cytotoxic cytokines and create a complex inflammatory milieu (Howell et al., 2011; Roberta Magliozzi et al., 2007). Whether a cell-mediated immune response, local release of cytotoxic compounds or environmental changes substantially shift the burden of repair from OPCs to surviving oligodendrocytes in human MS remains to be explored.

### Specificity of myelin repair

The regeneration of oligodendrocytes in different locations and the strong, cell intrinsic control of cell size appear to constrain where myelin sheaths are formed. However, when a regenerated oligodendrocyte had access to a territory that was previously myelinated, it was capable of establishing sheaths along axons that were previously myelinated, indicating that the local factors which initially influenced axon selection were retained after demyelination. In these regions, myelin sheaths were often reformed at a similar position along axons, even when sheaths were isolated from neighboring sheaths and therefore could have extended over a much larger area. Although previous studies suggest that components of the NOR are redistributed after demyelination in the PNS (England, Gamboni, Levinson, & Finger, 1990) and in MS lesions (Coman et al., 2006; Craner et al., 2004; Dupree et al., 2004), our studies reveal that βIV-spectrin, which forms a complex with Ankyrin-G to link voltage-gated sodium channels to the actin cytoskeleton at NORs (Susuki et al., 2016), remains clustered (without flanking myelin sheaths) with continued administration of cuprizone for up to six weeks (Figure 7D,F), suggesting that axonal guideposts remain for many weeks after demyelination. As these studies were performed using post-hoc immunostaining, we do not yet know if these βIV-spectrin puncta are located at previous nodal positions; future longitudinal studies using fluorescently tagged nodal components will help define the stability of NORs after demyelination. The time course over which these components are removed from demyelinated axons could influence the subsequent distribution of myelin, with prolonged demyelinating injuries leading to greater loss of remyelination specificity. Stabilizing the structural elements of previously myelinated axonal domains could therefore represent a potential target for remyelinating therapies.

Because territory overlap between original and regenerated oligodendrocytes was only ∼60%, considerable new axonal territory was accessed during the reparative period, leading to novel sheaths that altered the global pattern of cortical myelin. However, even in areas where the territories of regenerated and original oligodendrocytes overlapped, new myelin sheaths were often formed on portions of axons that were not previously myelinated. These findings highlight the probabilistic nature of myelination, which is influenced by many dynamic factors, such as neural activity, axon size and metabolic state that could alter target selection and stabilization of nascent sheaths (Klingseisen et al., 2019).

### Implications for myelin repair and cognitive function

Oligodendrocytes perform crucial roles in the CNS, enhancing the propagation of action potentials and reducing the metabolic cost to do so, providing metabolic support for axons far removed from their cell bodies and controlling excitability by influencing the distribution of voltage-gated channels and promoting clearance of extracellular potassium. Thus, the reorganization of myelin that occurs in cortical circuits during regeneration may have profound functional consequences on cognition and behavior. It will be important to determine whether these functional aspects of oligodendrocyte-neuron interactions are restored following remyelination. Our studies focused exclusively on regeneration in the somatosensory cortex of young adult mice. It is not yet known if these changes mimic regeneration in other cortical regions, or if the spatial and temporal aspects of myelin replacement in the cortex vary with age. Moreover, due to limitations in resolving sheaths in deeper layers of the cortex, we do not yet know whether regeneration deficits extend to layers IV-VI. It will also be critical to define the spatial and temporal changes in myelin sheath thickness, as this varies considerably between different axons and has been shown to be a substrate for plasticity in the cortex (Gibson et al., 2014).

Our analysis was restricted to discrete volumes in the cortical mantle. As we are not yet able to monitor myelination patterns along the entire length of individual axons, it is possible that the position of sheaths may change after regeneration, but that the overall myelin content along a given axon is conserved. Alternatively, from a functional standpoint, it may not be necessary to reform the precise pattern of myelination, as long as the relative amount of myelin along different classes of neurons is preserved. At present, our ability to predict the consequences of these changing myelin patterns and the spatial differences in oligodendrocyte regeneration are limited by our knowledge about the function of myelin in cortical gray matter. As recent studies suggest that even subtle changes in oligodendrogenesis can alter behavioral performance (McKenzie et al., 2014; Xiao et al., 2016), the impact of these changes may be profound. The ability to monitor the regeneration of myelin sheaths with high spatial and temporal resolution *in vivo* within defined circuits provides new opportunities to evaluate the effectiveness of potential therapeutic interventions (Deshmukh et al., 2013; Early et al., 2018; Mei et al., 2014; Rankin et al., 2019) and a platform to explore the functional consequences of myelin reorganization.

## MATERIALS AND METHODS

### Animal care and use

Female and male adult mice were used for experiments and randomly assigned to experimental groups. All mice were healthy and did not display any overt behavioral phenotypes, and no animals were excluded from the analysis. Generation and genotyping of BAC transgenic lines from *Mobp*-*EGFP* (Gensat) (Hughes et al., 2018) have been previously described. Mice were maintained on a 12-h light/dark cycle, housed in groups no larger than 5, and food and water were provided *ad libitum* (except during cuprizone-administration, see below). All animal experiments were performed in strict accordance with protocols approved by the Animal Care and Use Committee at Johns Hopkins University.

### Cranial Windows

Cranial windows were prepared as previously described (Holtmaat et al., 2012; Hughes et al., 2018). Briefly, mice 7 to 10 weeks old were anesthetized with isoflurane (induction, 5%; maintenance, 1.5-2%, mixed with 0.5L/min O_2_), and their body temperature was maintained at 37° C with a thermostat-controlled heating plate. The skin over the right hemisphere was removed and the skull cleaned. A 2 × 2 or 3 × 3 mm region of skull over somatosensory cortex (- 1.5 mm posterior and 3.5 mm lateral from bregma) was removed using a high-speed dental drill. A piece of cover glass (VWR, No. 1) was placed in the craniotomy and sealed with VetBond (3M), then dental cement (C&B Metabond) and a custom metal plate with a central hole was attached to the skull for head stabilization.

### *In vivo* two photon microscopy

*In vivo* imaging sessions began 2 to 3 weeks after cranial window procedure (Baseline). After the baseline imaging session, mice were randomly assigned to cuprizone or control conditions. During imaging sessions, mice were anesthetized with isoflurane and immobilized by attaching the head plate to a custom stage. Images were collected using a Zeiss LSM 710 microscope equipped with a GaAsP detector using a mode-locked Ti:sapphire laser (Coherent Ultra) tuned to 920 nm. The average power at the sample during imaging was < 30 mW. Vascular landmarks were used to identify the same cortical area over longitudinal imaging sessions. Image stacks were 425 μm × 425 μm × 110 μm (2048 × 2048 pixels, corresponding to cortical layer I, Zeiss 20x objective), 425 μm × 425 μm × 550 μm (1024 × 1024 pixels) or 850 μm × 850 μm × 550 μm (1024 × 1024 pixels; corresponding to layers I – IV), relative the cortical surface. Mice were imaged every 1 to 7 days, for up to 15 weeks.

### Cuprizone administration

At 9 to 11 weeks of age, male and female *Mobp-EGFP* mice were fed a diet of milled, irradiated 18% protein rodent diet (Teklad Global) alone (control) or supplemented with 0.2% w/w bis(cyclohexanone) oxaldihydrazone (Cuprizone, Sigma-Aldrich) in custom gravity-fed food dispensers for 3 to 6 weeks. Both control and experimental condition mice were returned to regular pellet diet during the recovery period (Baxi et al., 2017).

### Immunohistochemistry

Mice were deeply anesthetized with sodium pentobarbital (100 mg/kg b.w.) and perfused transcardially with 4% paraformaldehyde (PFA in 0.1 M phosphate buffer, pH 7.4). Brains were then post-fixed in 4% PFA for 12 to 18 hours, depending on antibody sensitivity to fixation, before being transferred to a 30% sucrose solution (in PBS, pH 7.4). For horizontal sections, cortices were flat-mounted between glass slides and postfixed in 4% PFA for 6 to 12 hours at 4° C, transferred to 30% sucrose solution (in PBS, pH 7.4). Tissue was stored at 4° C for more than 48 h before sectioning. Brains were extracted, frozen in TissueTek, sectioned (−1.5 mm posterior and 3.5 mm lateral from bregma) at 30 to 50 μm thickness on a cryostat (Thermo Scientific Microm HM 550) at −20° C. Immunohistochemistry was performed on free-floating sections. Sections were preincubated in blocking solution (5% normal donkey serum, 0.3% Triton X-100 in PBS, pH 7.4) for 1 or 2 hours at room temperature, then incubated for 24 to 48 hours at 4° C or room temperature in primary antibody (listed in Key Resources Table). Secondary antibody (see Key Resources Table) incubation was performed at room temperature for 2 to 4 hours. Sections were mounted on slides with Aqua Polymount (Polysciences). Images were acquired using either an epifluorescence microscope (Zeiss Axio-imager M1) with Axiovision software (Zeiss) or a confocal laser-scanning microscope (Zeiss LSM 510 Meta; Zeiss LSM 710; Zeiss LSM 880). For glial cell counts, individual images of coronal sections were quantified by a blinded observer for number of NG2+, GFAP+ and GFP+ cells within a 425 μm × 500 μm region, and divided into 425 μm × 100 μm zones from the pial surface (Supplementary Figure 1). Immunostaining for nodal components was performed as above, except mice were transcardially perfused with PBS only and post-fixed in 4% PFA for 50 minutes.

### Image processing and analysis

Image stacks and time series were analyzed using FIJI/ImageJ. For presentation in figures, image brightness and contrast levels were adjusted for clarity. Myelin sheath images were additionally de-noised with a 3-D median filter (radius 0.5 to 1.5 pixels). Longitudinal image stacks were registered using FIJI plugin “Correct 3D Drift” (Parslow, Cardona, & Bryson-Richardson, 2014) and then randomized for analysis by a blinded observer.

### Cell tracking

Individual oligodendrocytes were followed in four dimensions using custom FIJI scripts (Hughes et al., 2018) or with SyGlass (IstoVisio) virtual reality software by defining individual EGFP+ cell bodies at each time point, recording *xyz* coordinates, and defining cellular behavior (new, lost, or stable cells). Oligodendrocytes that were characterized as “lost” initially lost EGFP signal in processes and myelin sheaths, before complete loss of signal from the cell body position. A “new” oligodendrocyte appeared with novel processes and internodes absent in baseline images. Dynamics of cell body positions were analyzed with custom MATLAB scripts, and cross-time point comparisons of 3-D coordinates were corrected by adding the average vector of movement of all cells between those timepoints (to account for imperfect image registration and expansion/contraction of brain volume over time). For quantification between different 100 μm depths, cells were binned between planes horizontal to the plane of the pia, and included cells were found by Delaunay triangulation.

### Myelin sheath analysis

Registered longitudinal *in vivo* Z-stacks collected from *Mobp-EGFP* mice were acquired using two-photon microscopy. Similar to that described previously (Hughes et al., 2018), all myelin sheaths within a volume of 100 μm × 100 μm × 100 μm from the pial surface were traced in FIJI using Simple Neurite Tracer (Longair, Baker, & Armstrong, 2011) at the baseline or final recovery time-point. Then, using registered time-series images from baseline to final recovery time-point, each myelin sheath was categorized as having 0, 1 or 2 myelin sheath neighbors (Figure 4), whether it was stable (connected via cytoplasmic process to same cell at baseline and at 5 weeks recovery), lost (present at baseline, but not at 5 weeks recovery), replaced (≥ 50% of the original sheath length was replaced by a sheath connected *via* cytoplasmic process to a regenerated oligodendrocyte), or novel (a sheath not present at baseline that was connected to a regenerated oligodendrocyte). If it was a stable or replaced myelin sheath, we determined whether the baseline myelin sheath had the same or different number of neighboring myelin sheaths. Myelin sheaths within the field that could not be definitively categorized were classified as “undefined”. Myelin paranodes were identified by increased EGFP fluorescence intensity (Hughes et al., 2018). Nodes of Ranvier were confirmed by plotting an intensity profile across the putative node; if the profile consisted of two local maxima separated by a minimum less than that of the internode, and the length of the gap between EGFP+ processes was < 5 μm, the structure was considered a node. For each field, myelin sheaths were traced by one investigator and independently assessed by a second investigator.

### Analysis of temporal and spatial dynamics of individual oligodendrocytes

Registered longitudinal *in vivo* Z-stacks collected from *Mobp-EGFP* mice were acquired using two-photon microscopy every 1-3 days to follow the dynamics of newly formed mature oligodendrocytes within cortical layer I at high resolution (200 - 400 μm *x-y*, 100 - 120 μm *z*; 2048 × 2048 pixels). Using images from day of appearance, all processes originating from the cell body, branch points, and individual myelin sheaths were traced in FIJI using Simple Neurite Tracer (Longair et al., 2011). Traced segments were put through a smoothing function prior to length calculations to reduce artifacts of jagged traces. The fate of each process and myelin sheath (stable, lost) and changes in length (stable, growth, retraction) were determined for up to 14 days per cell.

### Single oligodendrocyte territory analysis

3-D coordinates of traces from all sheaths of a single oligodendrocyte were averaged to find the center of mass (to reduce artifactual territory volume that would be above the pia resulting from centering at the cell body). From this center, ellipsoid volumes were calculated from *x-y* and *z* radii, working backwards from the distance of the furthest sheath voxel in 1-μm increments in each dimension. The ellipsoid dimensions for each traced cell was determined by the combination of *x-y* and *z* radii that produced the smallest volume containing ≥ 80% of the sheath voxels for that cell. Voxels within the ellipsoid were calculated by 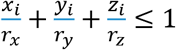 where (*x_i_*, *y_i_*, *z_i_*) is a voxel belonging to a sheath, and *r_x_*, *r_y_*, and *r_z_* are the radii being tested. Because all traced cell morphologies were not found to be significantly different from radial, the x and y radii were held equivalent. The average *x-y* and *z* radii across all traced baseline, control, or regenerated cells were used as standard dimensions for an oligodendrocyte “territory” in subsequent calculations.

Total territory volume for a single time-point (either baseline or 5 weeks recovery) was calculated in the following manner. In a 3-D matrix with dimensions, 425 μm × 425 μm × 33 μm corresponding to the number of voxels in the top zone of the imaged volume, all voxels contained within the territories of each cell were represented as 1’s, and all voxels outside of cell territories were represented as 0’s. Voxels within territories of more than one cell had values equal to the number of territories they were within. Total baseline volume “replaced” by regenerated territories was calculated as: ((5-week recovery matrix – baseline matrix) ≤ 0) + baseline matrix. The total proportion of baseline volume overlapped by regenerated territories was then calculated as: “replaced volume” / baseline volume. Because each imaged region had varying numbers of oligodendrocytes at baseline and regenerated oligodendrocytes in recovery, final territory overlap proportions were scaled in the following manner: total overlap × (number of cells @ baseline ÷ number of cells @ 5 weeks recovery).

### Analysis of nodal components

Following immunostaining, images were acquired at 63x on a Zeiss 880 microscope at high resolution (135 × 135 × 40 μm, 2048 × 2048 pixels). Images were processed in FIJI (background subtraction, rolling ball 60 pixels; 3-D median filter, 1.2 pixels XY 0.5 pixels Z). Images were acquired from sections immunostained for Caspr and βIV-spectrin, or both βIV-spectrin and Ankyrin-G. For images with both nodal components labeled, we found complete overlap of signal. In these cases, images were processed with “Image Calculator” in FIJI to reduce noise between channels, subtracting Ankyrin-G from βIV-spectrin signal. Images were imported into IMARIS and 3-D positions of all nodal signal and paranodal puncta (Caspr) were resolved with low- and high-pass filters and “Surface” and “Spot” functions. Axon initial segments were excluded using a size cutoff of ≤ 6 µm. Custom MATLAB scripts were used to detect proximity (nearest neighbor Euclidian distance, threshold 3.5 μm radius) of nodal puncta to Caspr puncta to determine proportions of nodes of Ranvier (2 Caspr puncta within 3.5 μm), heminodes (1 Caspr punctum within 3.5 μm), or isolated (no nearby Caspr puncta). Coordinates from one channel were rotated 90 degrees in the *x-y* plane before running the proximity analysis again to confirm observed proportions were not due to chance.

### Statistical analysis

No statistical tests were used to predetermine sample sizes, but our sample sizes are similar to those reported in previous publications (Hughes et al., 2018). Statistical analyses were performed with MATLAB (Mathworks) or Excel (Microsoft). Significance was typically determined using N-way ANOVA test with Bonferroni correction for multiple comparisons. Each figure legend otherwise contains the statistical tests used to measure significance and the corresponding significance level (p value). N represents the number of animals used in each experiment. Data are reported as mean ± SEM and p < 0.05 was considered statistically significant.

### Data availability

All published code, tools, and reagents will be shared on an unrestricted basis; requests should be directed to the corresponding authors.

## ACKNOWLEDGEMENTS

We thank Dr. M Pucak and N Ye for technical assistance, T Shelly for machining expertise, M Bhat for gift of the anti-Caspr antibody and members of the Bergles laboratory for discussions. J Orthmann-Murphy was supported by grants from the National Multiple Sclerosis Society and the Hilton Foundation. C Call and G Molina-Castro were supported by National Science Foundation Graduate Research Fellowships. Funding was provided by grants from the NIH (NS051509, NS050274, NS080153), a Collaborative Research Center Grant from the National Multiple Sclerosis Society, and the Dr. Miriam and Sheldon G Adelson Medical Research Foundation to D Bergles.

## SUPPLEMENTARY DATA

**Supplementary Figure 1:**
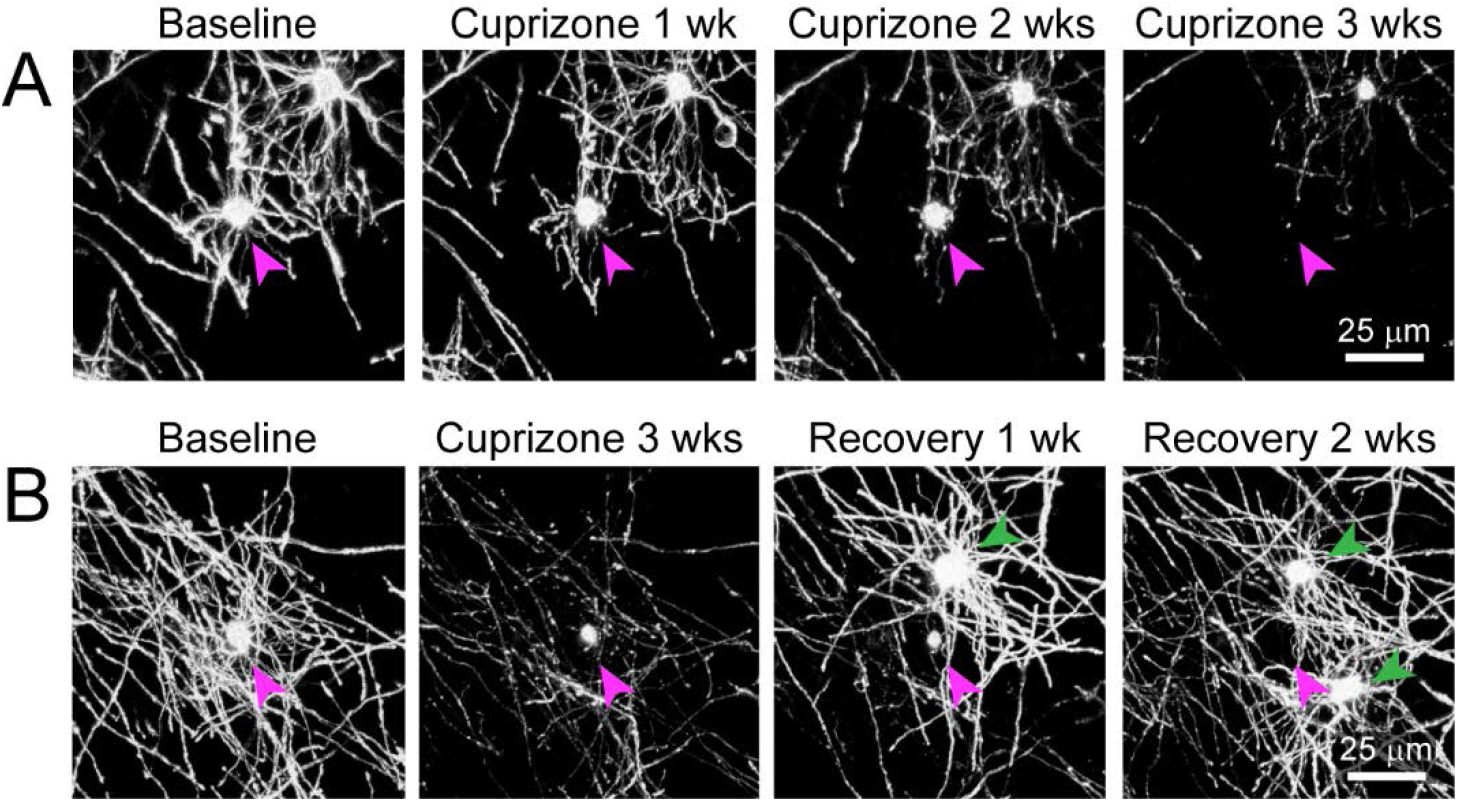
Degeneration of oligodendrocytes in cuprizone-treated *Mobp-EGFP* mice. Shown are two examples of individual oligodendrocytes tracked longitudinally using two photon *in vivo* imaging through chronic cranial windows in cuprizone-fed adult *Mobp-EGFP* mice. A: Example of an oligodendrocyte present at baseline (cell body denoted with magenta arrowhead) that loses GFP signal in processes and myelin sheaths and eventually cell body by 3 weeks of cuprizone treatment (maximum intensity projection of 156 μm × 156 μm × 45 μm volume). B. Example of an oligodendrocyte present at baseline that loses GFP signal in processes and myelin sheaths over a much longer time course than the cell in a, eventually disappearing at 3 weeks of recovery, after new oligodendrocytes (green arrowheads) are formed during recovery period (maximum intensity projection of 156 μm × 156 μm × 55 μm volume).

**Supplementary Figure 2:**
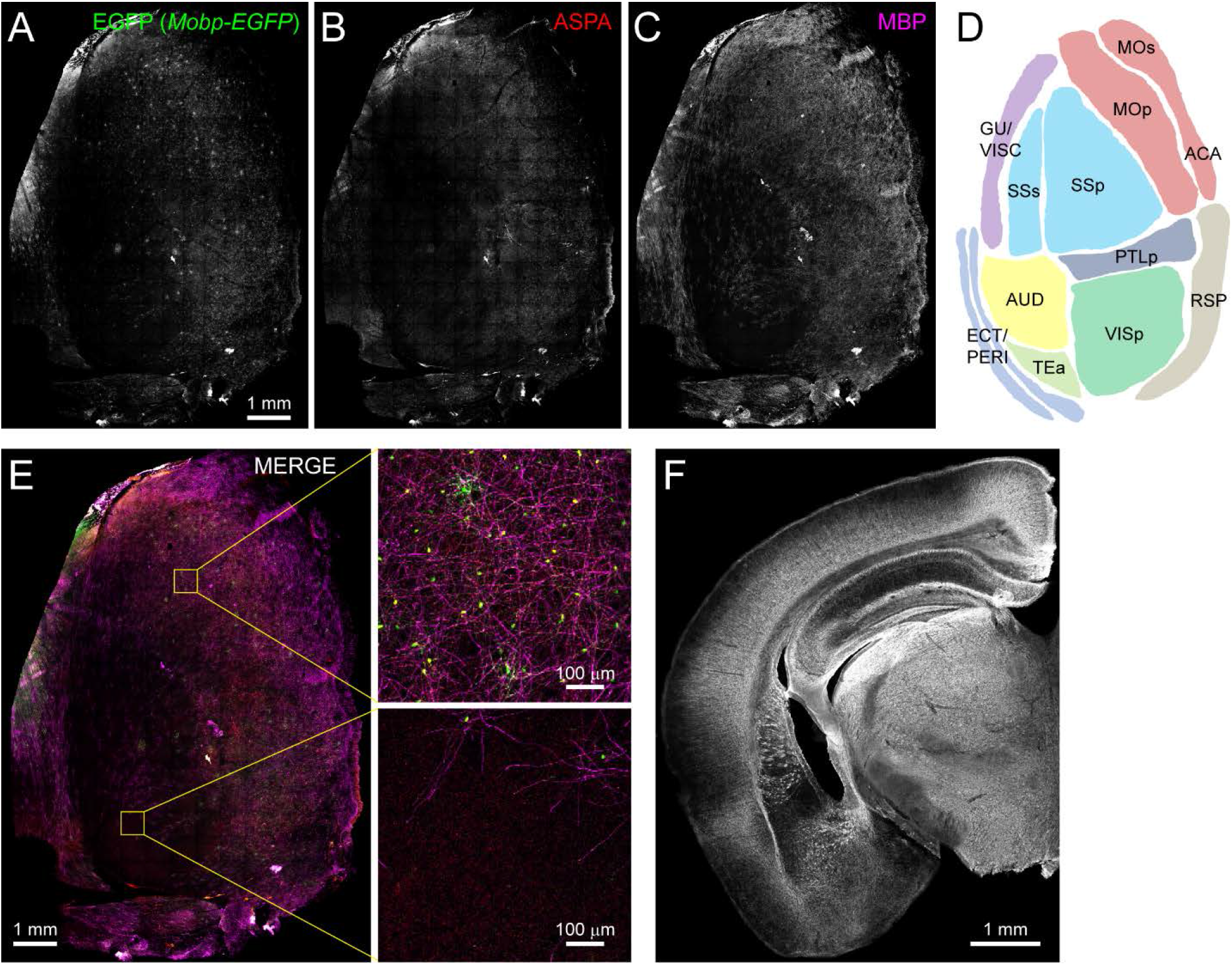
Myelin is not uniformly distributed throughout the adult rodent cortex. A-C: The left hemi-cortex from an adult *Mobp*-*EGFP* mouse is flattened and immunostained for EGFP (A), ASPA (B, oligodendrocytes) and MBP (C, myelin), merged together (E) and individual regions (schematized map in D) are expanded to illustrate that some cortical areas have a higher density of MBP+ myelin sheaths and GFP+/ASPA+ oligodendrocytes (top, primary somatomotor cortex) than others (bottom, auditory cortex). F: Coronal section from a 6-month old wild-type mouse immunostained for MBP shows regional heterogeneity of myelin across cortical mantle.

**Supplementary Figure 3:**
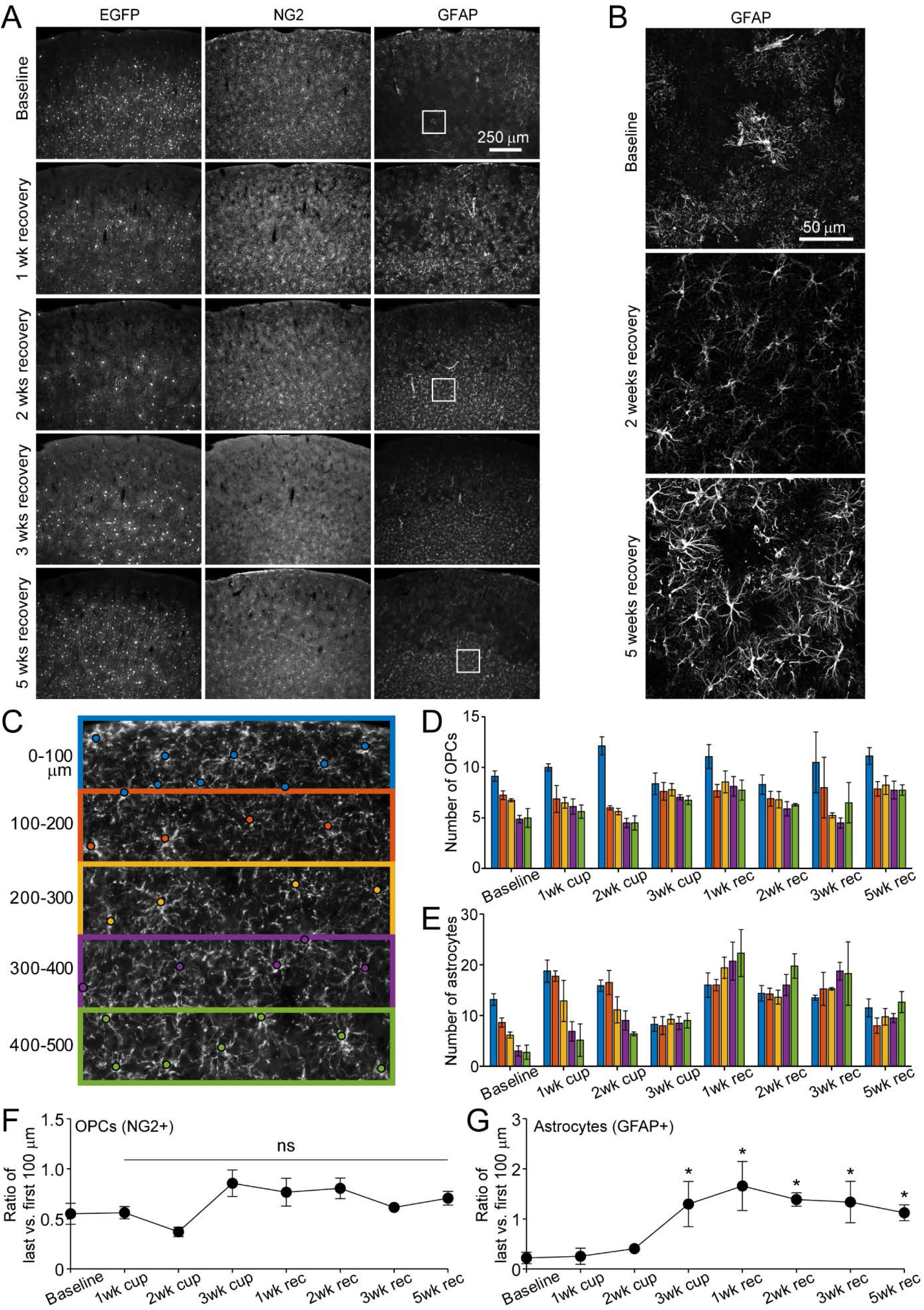
Distribution of astrocytes and oligodendrocyte precursor cells over the course of cuprizone treatment and recovery. A: Example coronal images from the brains of young adult *Mobp-EGFP* mice euthanized at baseline or following 3 weeks of cuprizone administration followed by 1, 2, 3 or 5 weeks recovery. Sections were immunostained for EGFP (oligodendrocytes), NG2 (OPCs) and GFAP (astrocytes). By 2 weeks of recovery, the relatively sparse distribution of EGFP+ cells represent newly formed cells (as demonstrated by *in vivo* imaging in Figure 1), with increasing number of new EGFP+ cells in lower cortical layers in later weeks of recovery. NG2+ OPC distribution remains constant over the course of damage and repair. GFAP+ cells increase in number after cuprizone and remain elevated in lower cortical regions over several weeks of recovery. B: Example maximum intensity projections of coronal sections from *Mobp-EGFP* mice euthanized at baseline, 2 weeks recovery or 5 weeks recovery, immunostained for GFAP (213 μm × 213 μm × 35 μm, from A). GFAP+ astrocyte morphology becomes reactive after exposure to cuprizone and maintains reactive morphology, even at 5 weeks of recovery. C: Example of a baseline somatosensory cortex coronal section from an *Mobp-EGFP* mouse, immunostained for NG2+, and divided into 100 μm zones from the pial surface to 500 μm in depth. The cell body location is marked with a circle. Each 100-μm zone color coded from pial surface corresponds bar colors in D and E. D-E: Quantification of cortical NG2+ cell distribution (D), and GFAP+ astrocytes (E) from brains of adult *Mobp-EGFP* mice euthanized at baseline (N = 4), 1 week of cuprizone (N = 4), 2 weeks of cuprizone (N = 4), 3 weeks of cuprizone (N = 4), 1 week of recovery (N = 8), 2 weeks of recovery (N = 5), or 3 weeks of recovery (N = 5 for NG2, N = 4 for GFAP), 5 weeks of recovery (N = 4). F-G. The ratio of cell number in the last (400-500 μm, green in C) versus first (0-100 μm, blue in C) zone for NG2+ OPCs (F) and GFAP+ astrocytes (G). Compared to baseline, the relative proportion of NG2 cells in top vs. bottom regions is stable over the course of cuprizone-treatment and recovery (F; p = 0.0858; Kruskal-Wallis one-way ANOVA with Fisher’s LSD) whereas the number of GFAP+ cells significantly increase in the bottom zone after 2 weeks of cuprizone and remain elevated (G, *p* = 0.0056, Kruskal-Wallis one-way ANOVA with Fisher’s LSD). Error bars are standard error of the mean.

**Supplementary Figure 4:**
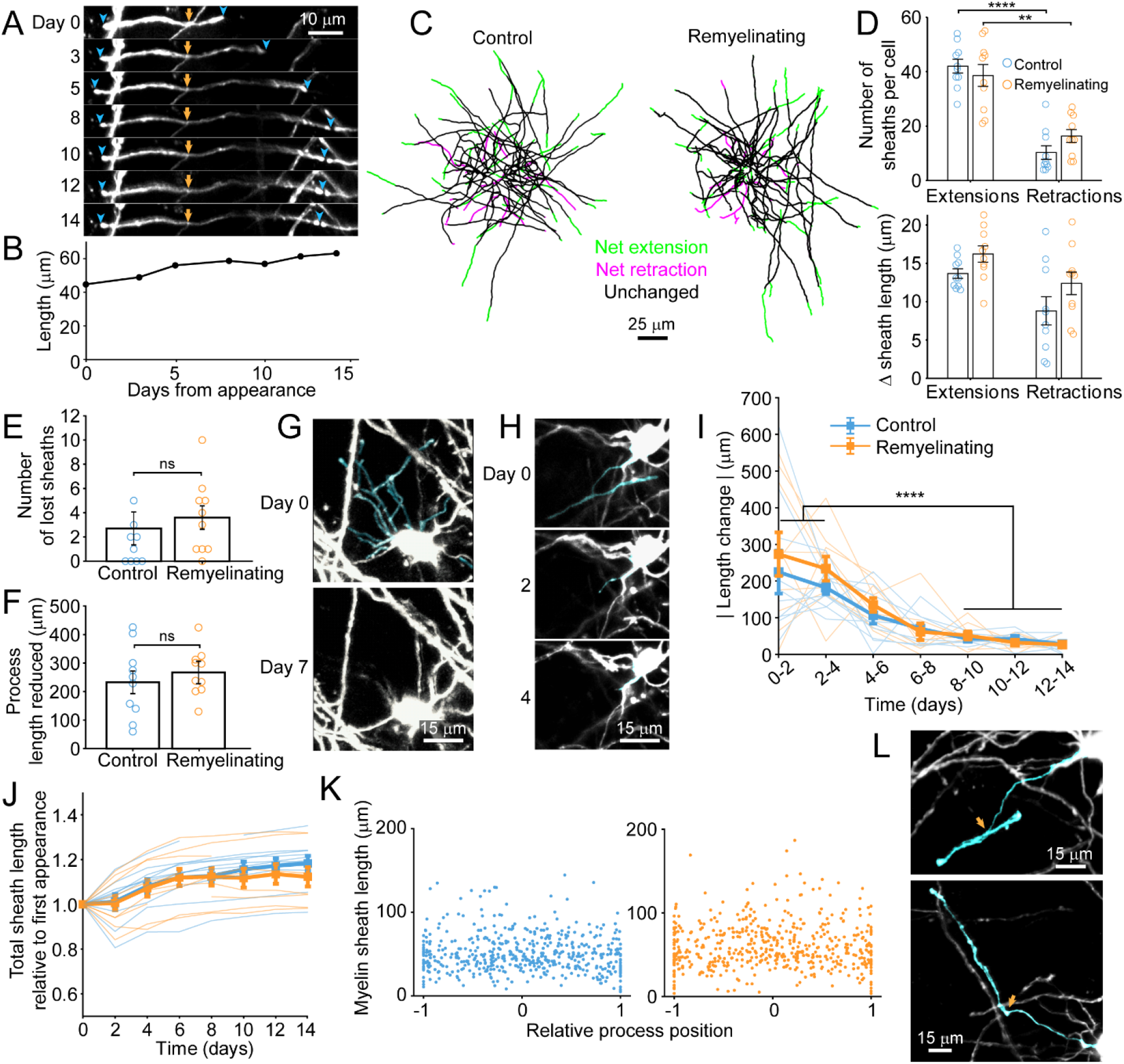
Dynamics of oligodendrocyte maturation in adult *Mobp*-*EGFP* mice. Every cell body, and associated myelin sheath belonging to newly appearing oligodendrocytes in cortical layer I were traced using Simple Neurite Tracer (Image J) on day of appearance and imaged every 1-3 days for up to 14 days. A: An individual myelin sheath (demarcated by blue arrowheads at the paranodal tips) at day 0 was followed over 14 days. The left-side paranode extends (through day 5) and then retracts and the right-side paranode extends to encounter a neighboring sheath and subsequently flanks a node of Ranvier (day 14). The yellow arrow marks the position where the oligodendrocyte process connects to the myelin sheath. B: The overall length of the individual sheath shown in A increases over 14 days. C: Example maximum intensity projections of traced processes and myelin sheaths from newly formed cells in control or cuprizone-treated mice on day of appearance, that were followed over 14 days and length of individual traced sheaths are denoted as either unchanged (black) or exhibiting net extension (green) or retraction (magenta). D: There were significantly more myelin sheaths undergoing extension than retraction in newly formed cells (control, p = 4.11 × 10^−7^; remyelinating, p = 0.00119, unpaired two-tailed t-tests, with Bonferroni correction for multiple comparisons) (top) but no difference in net length change of extensions or retractions (bottom), and no significant difference between control or remyelinating cells. E-F: There were low numbers of myelin sheaths lost in newly formed oligodendrocytes (E) and cytoplasmic processes were retracted (F); but there were no significant differences between control or remyelinating cells (sheaths, p = 0.907, unpaired two-tailed t-test; processes, p = 0.474, unpaired two-tailed t-test; control N = 10 cells, remyelinating N = 10 cells)). G-H: Examples of cytoplasmic processes and myelin sheaths (cyan) present at day 0 in a newly formed remyelinating cell, that are no longer present at day 7 (G) or day 4 (H). In H, the myelin sheath is dissolved first (day 2) and then the process connecting it to the cell body retracts completely by day 4. I-J: The absolute value of net total change in myelin sheath length over time is plotted for newly formed traced cells in control (blue) and cuprizone-treated (orange) cortex (thick line depicts means, thin lines represent individual cells). The majority of the length changes occur in the first 4 days after appearance (p = 2.28 × 10^−6^, N-way ANOVA with Bonferroni correction for multiple comparisons). J: The summed total length of all myelin sheaths per newly formed cell are plotted as a proportion of the total length at day of appearance. The overall trend is extension of myelin for both control and remyelinating cells. K: The length of individual myelin sheaths across all reconstructed cells plotted against the contact point of the cytoplasmic process to the myelin sheath, where 0 represents the center of an individual sheath (example in L, top panel, cyan process intersecting the cyan sheath towards the center of the sheath at orange arrow) and 1 or -1 are the distal tips of the sheath (example in L, bottom panel, cyan process intersecting at the paranode of the cyan sheath at orange arrow).

**Supplementary Video 1: Loss and replacement of oligodendrocytes.** Longitudinal *in vivo* imaging of demyelination and remyelination. This is a 392 μm × 392 μm × 100 μm volume shown as a maximum intensity projection that was repeatedly imaged through a chronic cranial window over the somatosensory cortex in an adult *Mobp-EGFP* mouse, at baseline, over 3 weeks of cuprizone administration, and then through 5 weeks of recovery. Scale bar is 50 μm.

**Link:** https://drive.google.com/file/d/18dTdmKwywMHbMt81p_64AshmRvgD8OEX/view?usp=sharing

**Supplementary Video 2: New oligodendrocytes are added in the upper cortical layers in adult mice.** Longitudinal imaging of an adult *Mobp-EGFP* mouse with a chronic cranial window fed sham diet. Region corresponds to images shown in Figure 1E, top row. Scale bar is 25 μm.

**Link:** https://drive.google.com/file/d/1b6up2_RWkh5qetrvYIqGv7kBQirTkFSi/view?usp=sharing

**Supplementary Video 3: Oligodendrocytes are lost and new cells appear after cuprizone-treatment.** Longitudinal imaging of an adult *Mobp-EGFP* mouse with a chronic cranial window fed 3 weeks of a cuprizone-supplemented diet followed through 5 weeks of recovery. Region corresponds to images shown in Figure 1E, bottom row. Scale bar is 25 μm.

**Link:** https://drive.google.com/file/d/1keWRJVo8kbN1khxA0tzdQzZV8GsPrL0j/view?usp=sharing

**Supplementary Video 4: Myelin sheaths are lost and novel sheaths are formed after cuprizone-treatment.** Longitudinal imaging of an adult *Mobp-EGFP* mouse with a chronic cranial window fed 3 weeks of a cuprizone-supplemented diet followed through 5 weeks of recovery. A myelin sheath at baseline (traced and pseudocolored magenta, overlayed in maximum intensity projection of longitudinally-imaged region), degenerates over time (only the traced sheath from baseline is shown in subsequent time-points, and is lost by 1 week of recovery). At 5 days of recovery a novel isolated sheath (not present at baseline, traced and pseudocolored in green in the 5 week recovery time-point overlay) appears, formed by a remyelinating oligodendrocyte not present at baseline (cell in 5 week recovery time-point overlay). Scale bar is 15 μm.

**Link:** https://drive.google.com/file/d/1fyVMqlAPOTH45SlOfgpWOg_yC-Pol45U/view?usp=sharing

**Supplementary Video 5: Isolated myelin sheaths are replaced.** Longitudinal imaging of an adult *Mobp-EGFP* mouse with a chronic cranial window fed 3 weeks of a cuprizone-supplemented diet followed through 5 weeks of recovery. An isolated myelin sheath at baseline (traced and pseudocolored magenta, overlayed in maximum intensity projection of longitudinally-imaged region), degenerates over time (only the traced sheath from baseline is shown in subsequent time-points, and is lost by 3 days of recovery). At 5 days of recovery a replacement isolated sheath appears (traced and pseudocolored in the 5 week recovery time-point overlay), formed by a remyelinating oligodendrocyte not present at baseline (cell in 5 week recovery time-point overlay). Scale bar is 15 μm.

**Link:** https://drive.google.com/file/d/1T6-Y0-rz7Gx32zjJMSaLcq9B6AAYETvM/view?usp=sharing

**Supplementary Video 6: Neighboring myelin sheaths with at least one neighbor are replaced and form a node of Ranvier in close proximity to one present at baseline.** Longitudinal imaging of an adult *Mobp-EGFP* mouse with a chronic cranial window fed 3 weeks of a cuprizone-supplemented diet followed through 5 weeks of recovery. Two neighboring myelin sheaths (pseudocolored magenta in baseline time-point), flank an unlabeled node of Ranvier (paranodal loops of myelin accumulate cytoplasmic EGFP in *Mopb-EGFP* mice (Hughes et al., 2018)); these sheaths were traced over each imaging time-points, and degenerate after cuprizone-treatment. Remyelinating oligodendrocytes (one shown in 5 week recovery time-point) form replacement myelin sheaths (pseudocolored green in 5 week recovery time-point); appear to flank an unlabeled node of Ranvier in a similar position as baseline. Scale bar is 15 μm.

**Link:** https://drive.google.com/file/d/1jA_lK-Cj3_TGlNfeJ32n-6GQ8E2OeLSv/view?usp=sharing

### Supplementary Table: Key Resources

#### Primary antibodies

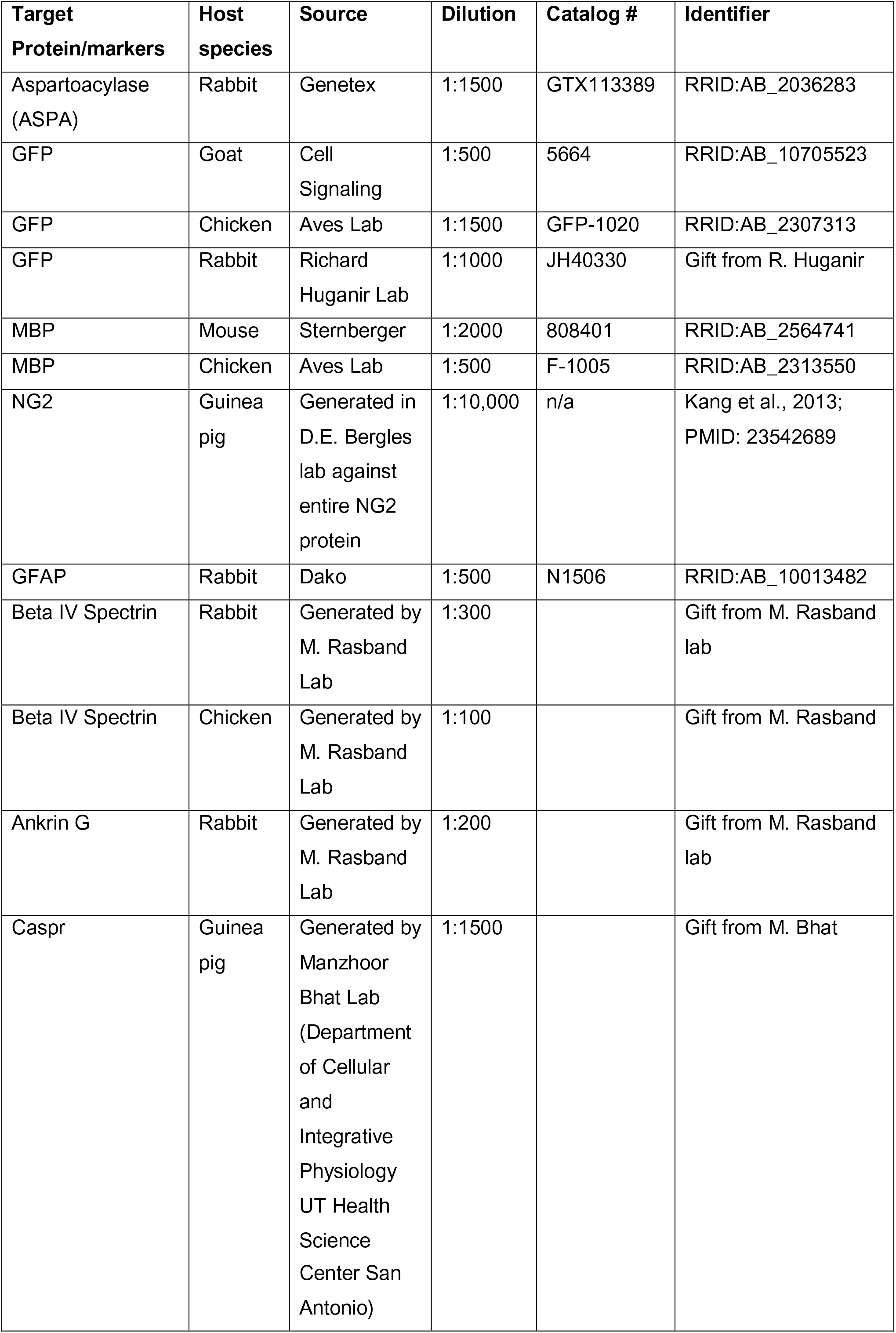

#### Secondary antibodies

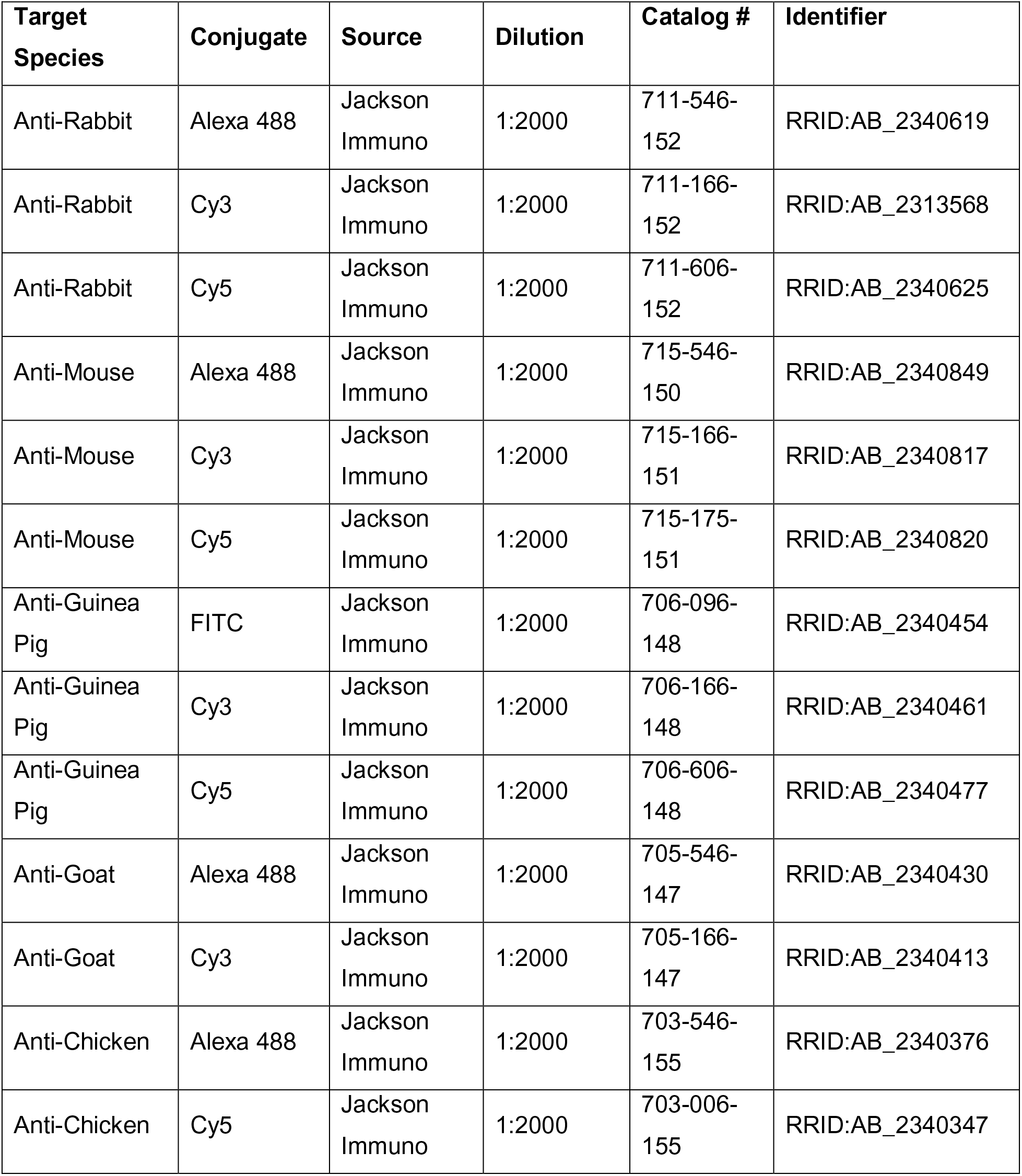

| Target Species | Conjugate | Source | Dilution | Catalog # | Identifier |
| --- | --- | --- | --- | --- | --- |
| Anti-Rabbit | Alexa 488 | Jackson Immuno | 1:2000 | 711-546-152 | RRID:AB_2340619 |
| Anti-Rabbit | Cy3 | Jackson Immuno | 1:2000 | 711-166-152 | RRID:AB_2313568 |
| Anti-Rabbit | Cy5 | Jackson Immuno | 1:2000 | 711-606-152 | RRID:AB_2340625 |
| Anti-Mouse | Alexa 488 | Jackson Immuno | 1:2000 | 715-546-150 | RRID:AB_2340849 |
| Anti-Mouse | Cy3 | Jackson Immuno | 1:2000 | 715-166-151 | RRID:AB_2340817 |
| Anti-Mouse | Cy5 | Jackson Immuno | 1:2000 | 715-175-151 | RRID:AB_2340820 |
| Anti-Guinea Pig | FITC | Jackson Immuno | 1:2000 | 706-096-148 | RRID:AB_2340454 |
| Anti-Guinea Pig | Cy3 | Jackson Immuno | 1:2000 | 706-166-148 | RRID:AB_2340461 |
| Anti-Guinea Pig | Cy5 | Jackson Immuno | 1:2000 | 706-606-148 | RRID:AB_2340477 |
| Anti-Goat | Alexa 488 | Jackson Immuno | 1:2000 | 705-546-147 | RRID:AB_2340430 |
| Anti-Goat | Cy3 | Jackson Immuno | 1:2000 | 705-166-147 | RRID:AB_2340413 |
| Anti-Chicken | Alexa 488 | Jackson Immuno | 1:2000 | 703-546-155 | RRID:AB_2340376 |
| Anti-Chicken | Cy5 | Jackson Immuno | 1:2000 | 703-006-155 | RRID:AB_2340347 |

#### Software and Algorithms

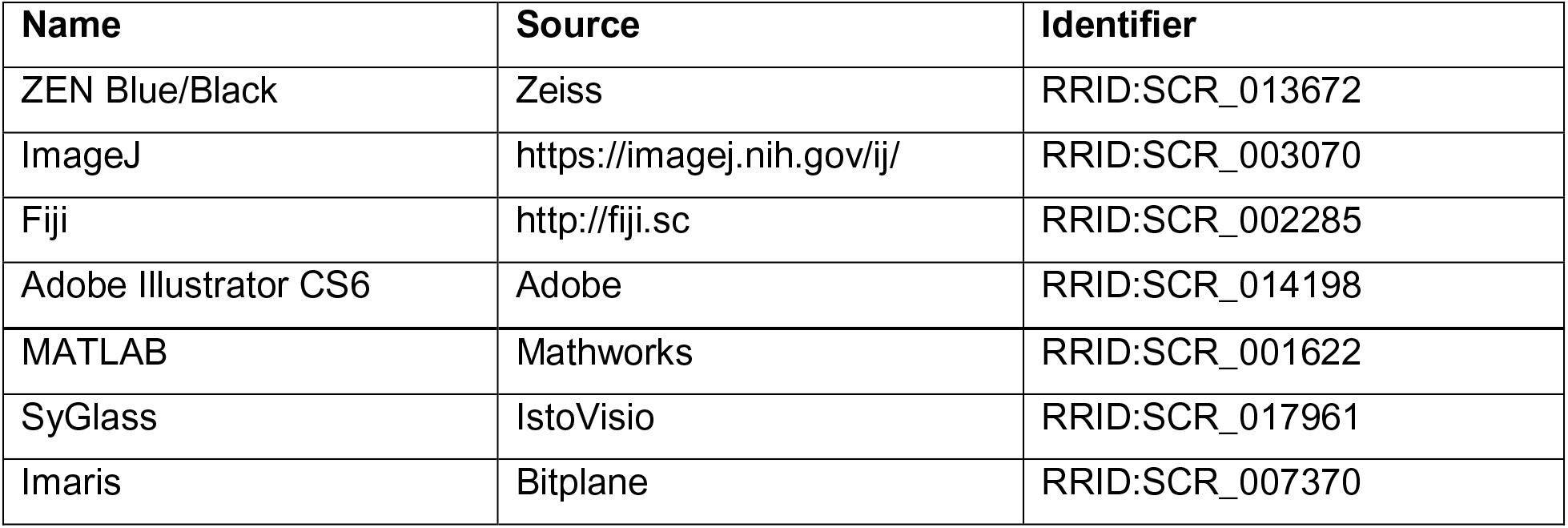

## REFERENCES

Araújo, S. E. S., Mendonça, H. R., Wheeler, N. A., Campello-Costa, P., Jacobs, K. M., Gomes, F. C. A., … Fuss, B. (2017). Inflammatory demyelination alters subcortical visual circuits. Journal of Neuroinflammation, 14(1), 162. https://doi.org/10.1186/s12974-017-0936-0

Auer, F., Vagionitis, S., & Czopka, T. (2018). Evidence for Myelin Sheath Remodeling in the CNS Revealed by In Vivo Imaging. Current Biology : CB, 28(4), 549–559.e3. https://doi.org/10.1016/j.cub.2018.01.017

Back, S. A., Tuohy, T. M. F., Chen, H., Wallingford, N., Craig, A., Struve, J., … Sherman, L. S. (2005). Hyaluronan accumulates in demyelinated lesions and inhibits oligodendrocyte progenitor maturation. Nature Medicine, 11(9), 966–972. https://doi.org/10.1038/nm1279

Baxi, E. G., DeBruin, J., Jin, J., Strasburger, H. J., Smith, M. D., Orthmann-Murphy, J. L., … Calabresi, P. A. (2017). Lineage tracing reveals dynamic changes in oligodendrocyte precursor cells following cuprizone-induced demyelination. GLIA, 65(12), 2087–2098. https://doi.org/10.1002/glia.23229

Baxi, E. G., DeBruin, J., Tosi, D. M., Grishkan, I. V., Smith, M. D., Kirby, L. A., … Gocke, A. R. (2015). Transfer of Myelin-Reactive Th17 Cells Impairs Endogenous Remyelination in the Central Nervous System of Cuprizone-Fed Mice. Journal of Neuroscience, 35(22), 8626–8639. https://doi.org/10.1523/JNEUROSCI.3817-14.2015

Beck, E. S., Sati, P., Sethi, V., Kober, T., Dewey, B., Bhargava, P., … Reich, D. S. (2018). Improved Visualization of Cortical Lesions in Multiple Sclerosis Using 7T MP2RAGE. American Journal of Neuroradiology, 39(3), 459–466. https://doi.org/10.3174/ajnr.A5534

Biteau, B., Hochmuth, C. E., & Jasper, H. (2011). Maintaining Tissue Homeostasis: Dynamic Control of Somatic Stem Cell Activity. Cell Stem Cell, 9(5), 402–411. https://doi.org/10.1016/j.stem.2011.10.004

Bock, D. D., Lee, W.-C. A., Kerlin, A. M., Andermann, M. L., Hood, G., Wetzel, A. W., … Reid, R. C. (2011). Network anatomy and in vivo physiology of visual cortical neurons. Nature, 471(7337), 177–182. https://doi.org/10.1038/nature09802

Calabrese, M., Poretto, V., Favaretto, A., Alessio, S., Bernardi, V., Romualdi, C., … Gallo, P. (2012). Cortical lesion load associates with progression of disability in multiple sclerosis. Brain, 135(10), 2952–2961. https://doi.org/10.1093/brain/aws246

Chang, A., Nishiyama, A., Peterson, J., Prineas, J., & Trapp, B. D. (2000). NG2-positive oligodendrocyte progenitor cells in adult human brain and multiple sclerosis lesions. The Journal of Neuroscience : The Official Journal of the Society for Neuroscience, 20(17), 6404–6412. https://doi.org/10.1523/JNEUROSCI.20-17-06404.2000

Chang, A., Staugaitis, S. M., Dutta, R., Batt, C. E., Easley, K. E., Chomyk, A. M., … Trapp, B. D. (2012). Cortical remyelination: A new target for repair therapies in multiple sclerosis. Annals of Neurology, 72(6), 918–926. https://doi.org/10.1002/ana.23693

Chong, S. Y. C., Rosenberg, S. S., Fancy, S. P. J., Zhao, C., Shen, Y.-A. A., Hahn, A. T., … Chan, J. R. (2012). Neurite outgrowth inhibitor Nogo-A establishes spatial segregation and extent of oligodendrocyte myelination. Proceedings of the National Academy of Sciences of the United States of America, 109(4), 1299–1304. https://doi.org/10.1073/pnas.1113540109

Coman, I., Aigrot, M. S., Seilhean, D., Reynolds, R., Girault, J. A., Zalc, B., & Lubetzki, C. (2006). Nodal, paranodal and juxtaparanodal axonal proteins during demyelination and remyelination in multiple sclerosis. Brain : A Journal of Neurology, 129(Pt 12), 3186–3195. https://doi.org/10.1093/brain/awl144

Craner, M. J., Newcombe, J., Black, J. A., Hartle, C., Cuzner, M. L., & Waxman, S. G. (2004). Molecular changes in neurons in multiple sclerosis: Altered axonal expression of Nav1.2 and Nav1.6 sodium channels and Na+/Ca2+ exchanger. Proceedings of the National Academy of Sciences, 101(21), 8168–8173. https://doi.org/10.1073/pnas.0402765101

Czopka, T., Ffrench-Constant, C., & Lyons, D. A. (2013). Individual oligodendrocytes have only a few hours in which to generate new myelin sheaths in vivo. Developmental Cell, 25(6), 599–609. https://doi.org/10.1016/j.devcel.2013.05.013

Deshmukh, V. A., Tardif, V., Lyssiotis, C. A., Green, C. C., Kerman, B., Kim, H. J., … Lairson, L. L. (2013). A regenerative approach to the treatment of multiple sclerosis. Nature, 502(7471), 327–332. https://doi.org/10.1038/nature12647

Dimou, L., Simon, C., Kirchhoff, F., Takebayashi, H., & Götz, M. (2008). Progeny of Olig2-expressing progenitors in the gray and white matter of the adult mouse cerebral cortex. The Journal of Neuroscience : The Official Journal of the Society for Neuroscience, 28(41), 10434–10442. https://doi.org/10.1523/JNEUROSCI.2831-08.2008

Dupree, J. L., Mason, J. L., Marcus, J. R., Stull, M., Levinson, R., Matsushima, G. K., & Popko, B. (2004). Oligodendrocytes assist in the maintenance of sodium channel clusters independent of the myelin sheath. Neuron Glia Biology, 1(3), 179–192. https://doi.org/10.1017/S1740925X04000304

Dutta, D. J., Woo, D. H., Lee, P. R., Pajevic, S., Bukalo, O., Huffman, W. C., … Fields, R. D. (2018). Regulation of myelin structure and conduction velocity by perinodal astrocytes. Proceedings of the National Academy of Sciences of the United States of America, 115(46), 11832–11837. https://doi.org/10.1073/pnas.1811013115

Early, J. J., Cole, K. L., Williamson, J. M., Swire, M., Kamadurai, H., Muskavitch, M., & Lyons, D. A. (2018). An automated high-resolution in vivo screen in zebrafish to identify chemical regulators of myelination. ELife, 7. https://doi.org/10.7554/eLife.35136

England, J. D., Gamboni, F., Levinson, S. R., & Finger, T. E. (1990). Changed distribution of sodium channels along demyelinated axons. Proceedings of the National Academy of Sciences of the United States of America, 87(17), 6777–6780. Retrieved from http://www.ncbi.nlm.nih.gov/pubmed/2168559

Filippi, M., Evangelou, N., Kangarlu, A., Inglese, M., Mainero, C., Horsfield, M. A., & Rocca, M. A. (2014). Ultra-high-field MR imaging in multiple sclerosis. Journal of Neurology, Neurosurgery, and Psychiatry, 85(1), 60–66. https://doi.org/10.1136/jnnp-2013-305246

Gibson, E. M., Purger, D., Mount, C. W., Goldstein, A. K., Lin, G. L., Wood, L. S., … Monje, M. (2014). Neuronal activity promotes oligodendrogenesis and adaptive myelination in the mammalian brain. Science (New York, N.Y.), 344(6183). https://doi.org/10.1126/science.1252304

Gouw, A. A., Seewann, A., Vrenken, H., van der Flier, W. M., Rozemuller, J. M., Barkhof, F., … Geurts, J. J. G. (2008). Heterogeneity of white matter hyperintensities in Alzheimer’s disease: post-mortem quantitative MRI and neuropathology. Brain, 131(12), 3286–3298. https://doi.org/10.1093/brain/awn265

Gudi, V., Moharregh-Khiabani, D., Skripuletz, T., Koutsoudaki, P. N., Kotsiari, A., Skuljec, J., … Stangel, M. (2009). Regional differences between grey and white matter in cuprizone induced demyelination. Brain Research, 1283, 127–138. https://doi.org/10.1016/j.brainres.2009.06.005

Herranz, E., Louapre, C., Treaba, C. A., Govindarajan, S. T., Ouellette, R., Mangeat, G., … Mainero, C. (2019). Profiles of cortical inflammation in multiple sclerosis by 11C-PBR28 MR-PET and 7 Tesla imaging. Multiple Sclerosis (Houndmills, Basingstoke, England), 1352458519867320. https://doi.org/10.1177/1352458519867320

Hill, R. A., Li, A. M., & Grutzendler, J. (2018). Lifelong cortical myelin plasticity and age-related degeneration in the live mammalian brain. Nature Neuroscience, 21(5), 683–695. https://doi.org/10.1038/s41593-018-0120-6

Holtmaat, A., de Paola, V., Wilbrecht, L., Trachtenberg, J. T., Svoboda, K., & Portera-Cailliau, C. (2012). Imaging neocortical neurons through a chronic cranial window. Cold Spring Harbor Protocols, 2012(6), 694–701. https://doi.org/10.1101/pdb.prot069617

Howell, O. W., Reeves, C. A., Nicholas, R., Carassiti, D., Radotra, B., Gentleman, S. M., … Reynolds, R. (2011). Meningeal inflammation is widespread and linked to cortical pathology in multiple sclerosis. Brain, 134(9), 2755–2771. https://doi.org/10.1093/brain/awr182

Hughes, E. G., Kang, S. H., Fukaya, M., & Bergles, D. E. (2013). Oligodendrocyte progenitors balance growth with self-repulsion to achieve homeostasis in the adult brain. Nature Neuroscience, 16(6), 668–676. https://doi.org/10.1038/nn.3390

Hughes, E. G., Orthmann-Murphy, J. L., Langseth, A. J., & Bergles, D. E. (2018). Myelin remodeling through experience-dependent oligodendrogenesis in the adult somatosensory cortex. Nature Neuroscience, 21(5), 696–706. https://doi.org/10.1038/s41593-018-0121-5

Ihara, M., Polvikoski, T. M., Hall, R., Slade, J. Y., Perry, R. H., Oakley, A. E., … Kalaria, R. N. (2010). Quantification of myelin loss in frontal lobe white matter in vascular dementia, Alzheimer’s disease, and dementia with Lewy bodies. Acta Neuropathologica, 119(5), 579–589. https://doi.org/10.1007/s00401-009-0635-8

Jeffery, N. D., & Blakemore, W. F. (1995). Remyelination of mouse spinal cord axons demyelinated by local injection of lysolecithin. Journal of Neurocytology, 24(10), 775–781. https://doi.org/10.1007/bf01191213

Kang, S. H., Fukaya, M., Yang, J. K., Rothstein, J. D., & Bergles, D. E. (2010). NG2+ CNS glial progenitors remain committed to the oligodendrocyte lineage in postnatal life and following neurodegeneration. Neuron, 68(4), 668–681.

Kidd, D., Barkhof, F., McConnell, R., Algra, P. R., Allen, I. V, & Revesz, T. (1999). Cortical lesions in multiple sclerosis. Brain : A Journal of Neurology, 122 (Pt 1), 17–26. https://doi.org/10.1093/brain/122.1.17

Kilsdonk, I. D., de Graaf, W. L., Soriano, A. L., Zwanenburg, J. J., Visser, F., Kuijer, J. P. A., … Wattjes, M. P. (2013). Multicontrast MR imaging at 7T in multiple sclerosis: highest lesion detection in cortical gray matter with 3D-FLAIR. AJNR. American Journal of Neuroradiology, 34(4), 791–796. https://doi.org/10.3174/ajnr.A3289

Kirby, L., Jin, J., Cardona, J. G., Smith, M. D., Martin, K. A., Wang, J., … Calabresi, P. A. (2019). Oligodendrocyte precursor cells present antigen and are cytotoxic targets in inflammatory demyelination. Nature Communications, 10(1), 3887. https://doi.org/10.1038/s41467-019-11638-3

Klingseisen, A., Ristoiu, A.-M., Kegel, L., Sherman, D. L., Rubio-Brotons, M., Almeida, R. G., … Lyons, D. A. (2019). Oligodendrocyte Neurofascin Independently Regulates Both Myelin Targeting and Sheath Growth in the CNS. Developmental Cell, 51(6), 730–744.e6. https://doi.org/10.1016/j.devcel.2019.10.016

Kotter, M. R., Li, W.-W., Zhao, C., & Franklin, R. J. M. (2006). Myelin Impairs CNS Remyelination by Inhibiting Oligodendrocyte Precursor Cell Differentiation. Journal of Neuroscience, 26(1), 328–332. https://doi.org/10.1523/JNEUROSCI.2615-05.2006

Kutzelnigg, A. (2005). Cortical demyelination and diffuse white matter injury in multiple sclerosis. Brain, 128(11), 2705–2712. https://doi.org/10.1093/brain/awh641

Lassmann, H., Brück, W., Lucchinetti, C., & Rodriguez, M. (1997). Remyelination in multiple sclerosis. Multiple Sclerosis Journal, 3(2), 133–136. https://doi.org/10.1177/135245859700300213

Lau, L. W., Keough, M. B., Haylock-Jacobs, S., Cua, R., Döring, A., Sloka, S., … Yong, V. W. (2012). Chondroitin sulfate proteoglycans in demyelinated lesions impair remyelination. Annals of Neurology, 72(3), 419–432. https://doi.org/10.1002/ana.23599

Lodato, S., & Arlotta, P. (2015). Generating neuronal diversity in the mammalian cerebral cortex. Annual Review of Cell and Developmental Biology, 31, 699–720. https://doi.org/10.1146/annurev-cellbio-100814-125353

Longair, M. H., Baker, D. A., & Armstrong, J. D. (2011). Simple Neurite Tracer: open source software for reconstruction, visualization and analysis of neuronal processes. Bioinformatics (Oxford, England), 27(17), 2453–2454. https://doi.org/10.1093/bioinformatics/btr390

Lucchinetti, C. F., Popescu, B. F. G., Bunyan, R. F., Moll, N. M., Roemer, S. F., Lassmann, H., … Ransohoff, R. M. (2011). Inflammatory cortical demyelination in early multiple sclerosis. The New England Journal of Medicine, 365(23), 2188–2197. https://doi.org/10.1056/NEJMoa1100648

Magliozzi, R., Reynolds, R., & Calabrese, M. (2018). MRI of cortical lesions and its use in studying their role in MS pathogenesis and disease course. Brain Pathology, 28(5), 735–742. https://doi.org/10.1111/bpa.12642

Magliozzi, Roberta, Howell, O., Vora, A., Serafini, B., Nicholas, R., Puopolo, M., … Aloisi, F. (2007). Meningeal B-cell follicles in secondary progressive multiple sclerosis associate with early onset of disease and severe cortical pathology. Brain : A Journal of Neurology, 130(Pt 4), 1089–1104. https://doi.org/10.1093/brain/awm038

Matsushima, G. K., & Morell, P. (2001). The neurotoxicant, cuprizone, as a model to study demyelination and remyelination in the central nervous system. Brain Pathology (Zurich, Switzerland), 11(1), 107–116. https://doi.org/10.1111/j.1750-3639.2001.tb00385.x

McKenzie, I. A., Ohayon, D., Li, H., de Faria, J. P., Emery, B., Tohyama, K., & Richardson, W. D. (2014). Motor skill learning requires active central myelination. Science (New York, N.Y.), 346(6207), 318–322. https://doi.org/10.1126/science.1254960

Mei, F., Fancy, S. P. J., Shen, Y.-A. A., Niu, J., Zhao, C., Presley, B., … Chan, J. R. (2014). Micropillar arrays as a high-throughput screening platform for therapeutics in multiple sclerosis. Nature Medicine, 20(8), 954–960. https://doi.org/10.1038/nm.3618

Mi, S., Miller, R. H., Lee, X., Scott, M. L., Shulag-Morskaya, S., Shao, Z., … Pepinsky, R. B. (2005). LINGO-1 negatively regulates myelination by oligodendrocytes. Nature Neuroscience, 8(6), 745–751. https://doi.org/10.1038/nn1460

Micheva, K. D., Wolman, D., Mensh, B. D., Pax, E., Buchanan, J., Smith, S. J., & Bock, D. D. (2016). A large fraction of neocortical myelin ensheathes axons of local inhibitory neurons. ELife, 5. https://doi.org/10.7554/eLife.15784

Miron, V. E., Boyd, A., Zhao, J.-W., Yuen, T. J., Ruckh, J. M., Shadrach, J. L., … Ffrench-Constant, C. (2013). M2 microglia and macrophages drive oligodendrocyte differentiation during CNS remyelination. Nature Neuroscience, 16(9), 1211–1218. https://doi.org/10.1038/nn.3469

Murtie, J. C., Macklin, W. B., & Corfas, G. (2007). Morphometric analysis of oligodendrocytes in the adult mouse frontal cortex. Journal of Neuroscience Research, 85(10), 2080–2086. https://doi.org/10.1002/jnr.21339

Neumann, B., Baror, R., Zhao, C., Segel, M., Dietmann, S., Rawji, K. S., … Franklin, R. J. M. (2019). Metformin Restores CNS Remyelination Capacity by Rejuvenating Aged Stem Cells. Cell Stem Cell, 25(4), 473–485.e8. https://doi.org/10.1016/j.stem.2019.08.015

Nicaise, A. M., Wagstaff, L. J., Willis, C. M., Paisie, C., Chandok, H., Robson, P., … Crocker, S. J. (2019). Cellular senescence in progenitor cells contributes to diminished remyelination potential in progressive multiple sclerosis. Proceedings of the National Academy of Sciences, 116(18), 9030–9039. https://doi.org/10.1073/pnas.1818348116

Nielsen, A. S., Kinkel, R. P., Madigan, N., Tinelli, E., Benner, T., & Mainero, C. (2013). Contribution of cortical lesion subtypes at 7T MRI to physical and cognitive performance in MS. Neurology, 81(7), 641–649. https://doi.org/10.1212/WNL.0b013e3182a08ce8

Oh, J., Ontaneda, D., Azevedo, C., Klawiter, E. C., Absinta, M., Arnold, D. L., … North American Imaging in Multiple Sclerosis Cooperative. (2019). Imaging outcome measures of neuroprotection and repair in MS. Neurology, 92(11), 10.1212/WNL.0000000000007099. https://doi.org/10.1212/WNL.0000000000007099

Parslow, A., Cardona, A., & Bryson-Richardson, R. J. (2014). Sample drift correction following 4D confocal time-lapse imaging. Journal of Visualized Experiments : JoVE, (86). https://doi.org/10.3791/51086

Peterson, J. W., Bö, L., Mörk, S., Chang, A., & Trapp, B. D. (2001). Transected neurites, apoptotic neurons, and reduced inflammation in cortical multiple sclerosis lesions. Annals of Neurology, 50(3), 389–400. Retrieved from http://www.ncbi.nlm.nih.gov/pubmed/11558796

Rankin, K. A., Mei, F., Kim, K., Shen, Y.-A. A., Mayoral, S. R., Desponts, C., … Bove, R. (2019). Selective Estrogen Receptor Modulators Enhance CNS Remyelination Independent of Estrogen Receptors. The Journal of Neuroscience : The Official Journal of the Society for Neuroscience, 39(12), 2184–2194. https://doi.org/10.1523/JNEUROSCI.1530-18.2019

Ruckh, J. M., Zhao, J.-W., Shadrach, J. L., van Wijngaarden, P., Rao, T. N., Wagers, A. J., & Franklin, R. J. M. (2012). Rejuvenation of regeneration in the aging central nervous system. Cell Stem Cell, 10(1), 96–103. https://doi.org/10.1016/j.stem.2011.11.019

Samanta, J., Grund, E. M., Silva, H. M., Lafaille, J. J., Fishell, G., & Salzer, J. L. (2015). Inhibition of Gli1 mobilizes endogenous neural stem cells for remyelination. Nature, 526(7573), 448–452. https://doi.org/10.1038/nature14957

Skripuletz, T., Lindner, M., Kotsiari, A., Garde, N., Fokuhl, J., Linsmeier, F., … Stangel, M. (2008). Cortical demyelination is prominent in the murine cuprizone model and is strain-dependent. The American Journal of Pathology, 172(4), 1053–1061. https://doi.org/10.2353/ajpath.2008.070850

Stedehouder, J., Couey, J. J., Brizee, D., Hosseini, B., Slotman, J. A., Dirven, C. M. F., … Kushner, S. A. (2017). Fast-spiking Parvalbumin Interneurons are Frequently Myelinated in the Cerebral Cortex of Mice and Humans. Cerebral Cortex (New York, N.Y.: 1991), 27(10), 5001–5013. https://doi.org/10.1093/cercor/bhx203

Stedehouder, J., & Kushner, S. A. (2017). Myelination of parvalbumin interneurons: a parsimonious locus of pathophysiological convergence in schizophrenia. Molecular Psychiatry, 22(1), 4–12. https://doi.org/10.1038/mp.2016.147

Su, Z., Yuan, Y., Chen, J., Zhu, Y., Qiu, Y., Zhu, F., … He, C. (2011). Reactive astrocytes inhibit the survival and differentiation of oligodendrocyte precursor cells by secreted TNF-α. Journal of Neurotrauma, 28(6), 1089–1100. https://doi.org/10.1089/neu.2010.1597

Susuki, K., Otani, Y., & Rasband, M. N. (2016). Submembranous cytoskeletons stabilize nodes of Ranvier. Experimental Neurology, 283, 446–451. https://doi.org/10.1016/J.EXPNEUROL.2015.11.012

Tomassy, G. S., Berger, D. R., Chen, H.-H., Kasthuri, N., Hayworth, K. J., Vercelli, A., … Arlotta, P. (2014). Distinct Profiles of Myelin Distribution Along Single Axons of Pyramidal Neurons in the Neocortex. Science, 344(6181), 319–324. https://doi.org/10.1126/science.1249766

Trapp, B. D., Nishiyama, A., Cheng, D., & Macklin, W. (1997). Differentiation and death of premyelinating oligodendrocytes in developing rodent brain. The Journal of Cell Biology, 137(2), 459–468. https://doi.org/10.1083/jcb.137.2.459

Vega-Riquer, J. M., Mendez-Victoriano, G., Morales-Luckie, R. A., & Gonzalez-Perez, O. (2019). Five Decades of Cuprizone, an Updated Model to Replicate Demyelinating Diseases. Current Neuropharmacology, 17(2), 129–141. https://doi.org/10.2174/1570159X15666170717120343

Walhovd, K. B., Johansen-Berg, H., & Káradóttir, R. T. (2014). Unraveling the secrets of white matter--bridging the gap between cellular, animal and human imaging studies. Neuroscience, 276(100), 2–13. https://doi.org/10.1016/j.neuroscience.2014.06.058

Werneburg, S., Jung, J., Kunjamma, R. B., Ha, S.-K., Luciano, N. J., Willis, C. M., … Schafer, D. P. (2020). Targeted Complement Inhibition at Synapses Prevents Microglial Synaptic Engulfment and Synapse Loss in Demyelinating Disease. Immunity, 52(1), 167–182.e7. https://doi.org/10.1016/j.immuni.2019.12.004

Xiao, L., Ohayon, D., McKenzie, I. A., Sinclair-Wilson, A., Wright, J. L., Fudge, A. D., … Richardson, W. D. (2016). Rapid production of new oligodendrocytes is required in the earliest stages of motor-skill learning. Nature Neuroscience, 19(9), 1210–1217. https://doi.org/10.1038/nn.4351

Yeung, M. S. Y., Djelloul, M., Steiner, E., Bernard, S., Salehpour, M., Possnert, G., … Frisén, J. (2019). Dynamics of oligodendrocyte generation in multiple sclerosis. Nature, 566(7745), 538–542. https://doi.org/10.1038/s41586-018-0842-3

Yeung, M. S. Y., Zdunek, S., Bergmann, O., Bernard, S., Salehpour, M., Alkass, K., … Frisén, J. (2014). Dynamics of oligodendrocyte generation and myelination in the human brain. Cell, 159(4), 766–774. https://doi.org/10.1016/j.cell.2014.10.011

Young, K. M., Psachoulia, K., Tripathi, R. B., Dunn, S.-J., Cossell, L., Attwell, D., … Richardson, W. D. (2013). Oligodendrocyte dynamics in the healthy adult CNS: evidence for myelin remodeling. Neuron, 77(5), 873–885. https://doi.org/10.1016/j.neuron.2013.01.006

Zhang, Y., Zhang, J., Navrazhina, K., Argaw, A. T., Zameer, A., Gurfein, B. T., … John, G. R. (2010). TGFbeta1 induces Jagged1 expression in astrocytes via ALK5 and Smad3 and regulates the balance between oligodendrocyte progenitor proliferation and differentiation. Glia, 58(8), 964–974. https://doi.org/10.1002/glia.20978

